# Loss of NR5A1 in Sertoli cells after sex determination changes their cellular identity and induces their death by anoikis

**DOI:** 10.1101/2023.01.04.522755

**Authors:** Sirine Souali-Crespo, Diana Condrea, Nadège Vernet, Betty Féret, Muriel Klopfenstein, Erwan Grandgirard, Violaine Alunni, Marie Cerciat, Matthieu Jung, Chloé Mayere, Serge Nef, Manuel Mark, Frédéric Chalmel, Norbert B. Ghyselinck

**Author notes:** These authors contributed equally to this work (Frédéric Chalmel, Norbert B. Ghyselinck).

## Abstract

NR5A1 is an orphan nuclear receptor crucial for gonadal development in mammals. In the mouse testis it is expressed both in Sertoli cells (SC) and Leydig cells (LC). To investigate its role posteriorly to sex determination, we have generated and analysed mice lacking NR5A1 in SC from embryonic day (E) 13.5 onwards (*Nr5a1*^SC−/−^ mutants). Ablation of *Nr5a1* impairs the expression of genes characteristic of SC identity (e.g., *Sox9, Amh*), makes SC to progressively die from E14.5 by a *Trp53*-independent mechanism, and induces disorganization of the testis cords, which, together, yields germ cells (GC) to prematurely enter meiosis and die, instead of becoming quiescent. Single-cell RNA-sequencing experiments revealed that *Nr5a1*-deficient SC acquire a pre-granulosa cell-like identity, and profoundly modify the landscape of the adhesion molecules and extracellular matrix they express. We propose therefore that SC lacking NR5A1 transdifferentiate and die by anoikis. Fetal LC do not display major changes in their transcriptome, indicating that SC are not required beyond E14.5 for their emergence or maintenance. In contrast, adult LC were missing in *Nr5a1*^SC−/−^ postnatal testes. In addition, adult males display Müllerian duct derivatives (i.e., uterus, vagina), as well as a decreased anogenital distance and a shorter penis that can be explained by loss of AMH production and defective HSD17B1- and HSD17B3-mediated synthesis of testosterone in SC during fetal life. Together, our findings indicate that *Nr5a1* expressed in SC after the period of sex determination safeguards SC identity, which maintains proper seminiferous cord organization and prevents GC to enter meiosis.

## INTRODUCTION

Testis and ovary both originate from a bipotential gonad, which is composed of somatic progenitor cells that are competent to adopt one or the other sex-specific cell fate, and of primordial germ cells (GC). Sex determination initiates at around embryonic day (E) 11.5, from when somatic progenitor cells undergo sex-specific cell differentiation. The specification of the supporting cell lineage [i.e., either Sertoli cells (SC) in the testis upon SRY expression, or pre-granulosa cells (pGrC) in the ovary upon WNT/CTNNB1 signalling stabilization] is the first essential step in this process. Following supporting cell differentiation, the sex-specific fate decision propagates to the GC and to other somatic lineages, including the steroidogenic cells Leydig cells (LC) in the testis, which, in turn, drive acquisition of primary and secondary sexual characters at later stages of development, through hormone secretion [1–3]. In humans, mutations in genes driving this complex developmental process can give rise to a group of defects known as disorders of sex development (DSD), the diagnosis and the management of which requires an understanding of the molecular mechanisms underpinning cell differentiation [4].

Amongst the genes responsible for DSD is *Nr5a1*, encoding the orphan nuclear receptor NR5A1 [5], a transcription factor expressed at early stages of development, when cells are partitioned into adrenal and gonadal primordium [6]. NR5A1 controls the expression of genes implicated in proliferation, survival and differentiation of somatic progenitor cells [7]. Therefore, knock-out of *Nr5a1* in the mouse results in regression of the gonads by E11.5 due to apoptosis of somatic cells, and agenesis of the adrenal gland [8]. From E11.5, NR5A1 and SRY are expressed in the bipotential supporting cells of male embryos, where they upregulate SOX9 expression, initiating thereby pre-SC and ultimately SC differentiation [2,3,9]. Once specified, the SC start forming cords by enclosing GC as early as E12.5 in the mouse [10], providing them with a specialized environment that promotes their survival and orchestrates their differentiation [11]. SC also (i) allow the differentiation of fetal LC (FLC) through paracrine Desert Hedgehog (DHH) signalling [12], (ii) produce, under the control of NR5A1 and SOX9, high levels of anti-Müllerian hormone (AMH), which triggers the regression of the Müllerian duct normally giving rise to the female genitalia [13].

Because gonads are absent in *Nr5a1*-null mice [8], studying the cell-specific roles of NR5A1 in the developing testis required analysis of mice bearing tissue-targeted mutations. Here we have analysed the outcome of *Nr5a1* ablation in SC from E13.5 onwards. We show that loss of NR5A1 from this stage induces some SC to transdifferentiate into female somatic cells, and to die by a *Trp53*-independent mechanism related to anoikis.

## RESULTS

### Deletion of *Nr5a1* in Sertoli cells

To achieve *Nr5a1* ablation in SC after sex determination, we introduced the *Plekha5*^Tg(AMH-cre)1Flor^ transgene [14] in mice bearing *lox*P-flanked alleles of *Nr5a1* and a Cre-dependent reporter transgene [15]. Cre-mediated recombination was assessed by immunohistochemistry (IHC) using antibodies recognizing the yellow fluorescent protein (YFP). Surprisingly, YFP was detected in all SC of embryonic day (E) 12.5 gonads (arrows, **Fig.1A,B**), indicating Cre-mediated excision occurred earlier than anticipated [14]. However, the NR5A1 protein was still detected in SC at E12.5 (**Fig.1A,B**). At E13.5, the nuclei of SC were all NR5A1-positive in the control (**Fig.1C**), but NR5A1-negative in the mutant testis (**Fig.1D**). Importantly, NR5A1 was still detected in the nuclei of LC, identified by their expression of the steroid 3 beta-hydroxy-steroid dehydrogenase type 1 (HSD3B1), in both control and mutant testes (**Fig.1E,F**). Thus, NR5A1 was lost selectively in SC from E13.5 onwards, generating males hereafter called *Nr5a1*^SC−/−^ mutants.

**Figure 1.**
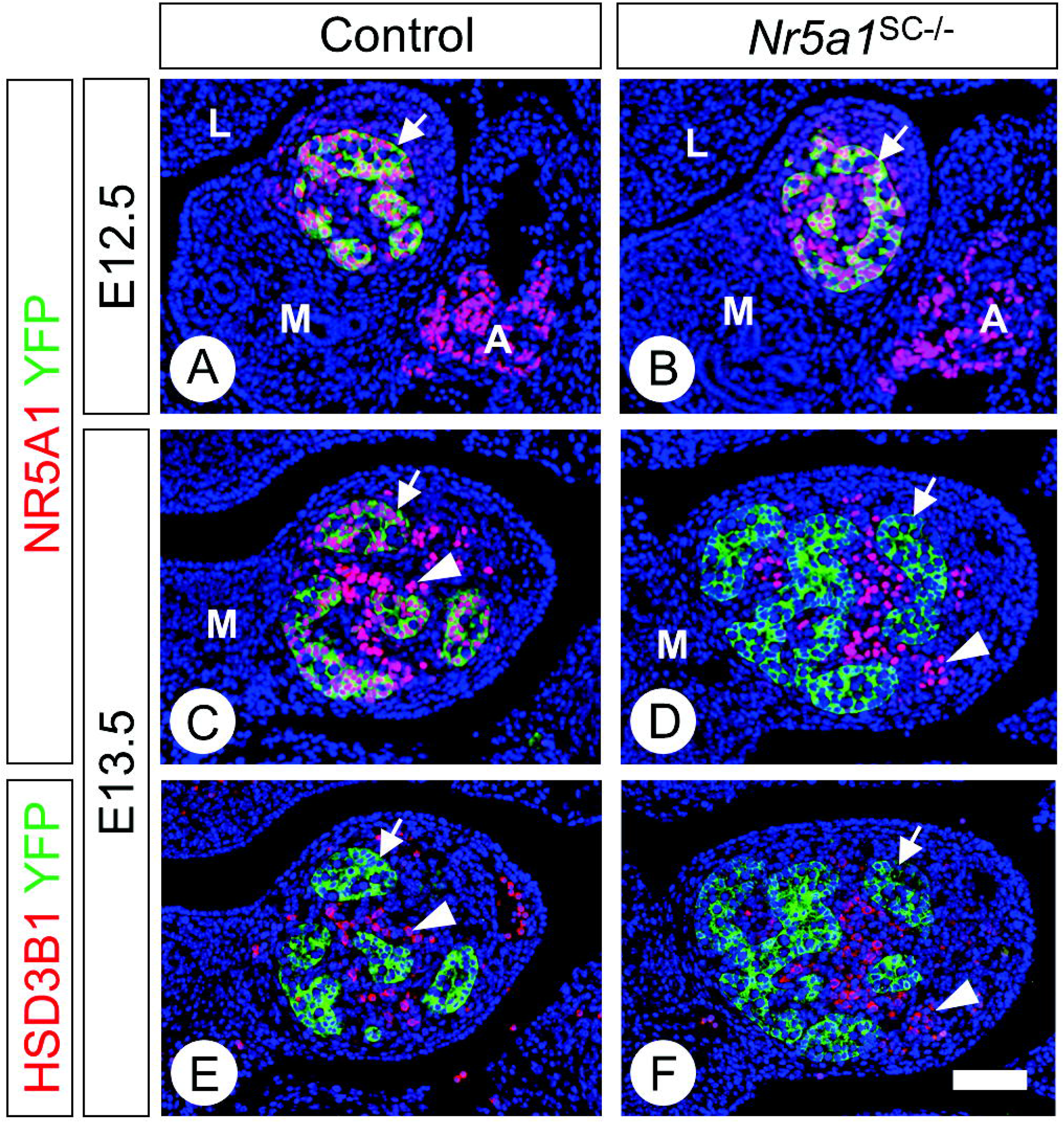
Ablation of *Nr5a1* is efficient from E13.5 onwards. **(A-F)** Detection of NR5A1 (red nuclear signals), YFP (green cytoplasmic signal), and HSD3B1 (red cytoplasmic signal) by IHC on transverse histological sections of the testis of a control (A,C,E) and a *Nr5a1*^SC−/−^ mutant fetus (B,D,F) at E12.5 (A,B) and E13.5 (C-F). Efficient excision of the reporter transgene by cre is assessed by YFP expression in virtually all SC (arrows) as early as E12.5 (A,B). However, loss of NR5A1 in SC is only achieved at E13.5 (compare C with D). At this stage, expression of NR5A1 is maintained in Leydig cells (arrowheads in C,D), as identified on consecutive sections by their expression of HSD3B1 (E,F). Nuclei are counterstained with DAPI (blue signal). Scale bar (in F): 50 μm.

### Ablation of *Nr5a1* in SC impairs expression of SOX9, SOX8, SOX10 and AMH

As NR5A1 regulates the expression of *Amh* and *Sox9* genes [9,16], we tested expression of these two proteins by IHC. Both AMH and SOX9 were detected at normal levels at E12.5 (**Fig.2A,B,G,H**). Their expression decreased progressively between E13.5 and E14.5 in *Nr5a1*^SC−/−^ mutant testes (**Fig.2C-F,I-L**). At E14.5, only few *Nr5a1*-deficient SC were SOX9-positive, some of which with cytoplasmic SOX9 (compare insets, **Fig.2K,L**). The expression of SOX8 and SOX10 was also decreased in *Nr5a1*^SC−/−^ gonads (**Suppl.Fig.1**). Accordingly, the steady state levels of *Amh, Sox9* and *Sox8* mRNAs was decreased in *Nr5a1*^SC−/−^ whole gonads (**Fig.2S**). The level of prostaglandin D2 synthase (*Ptgds*) mRNA, another SOX9 target-gene [17], was also reduced in E14.5 *Nr5a1*^SC−/−^ testes. This finding may explain the cytoplasmic localization of SOX9 in some SC since prostaglandin D2, the end product of PTGDS, is required for SOX9 nuclear translocation [18].

**Figure 2.**
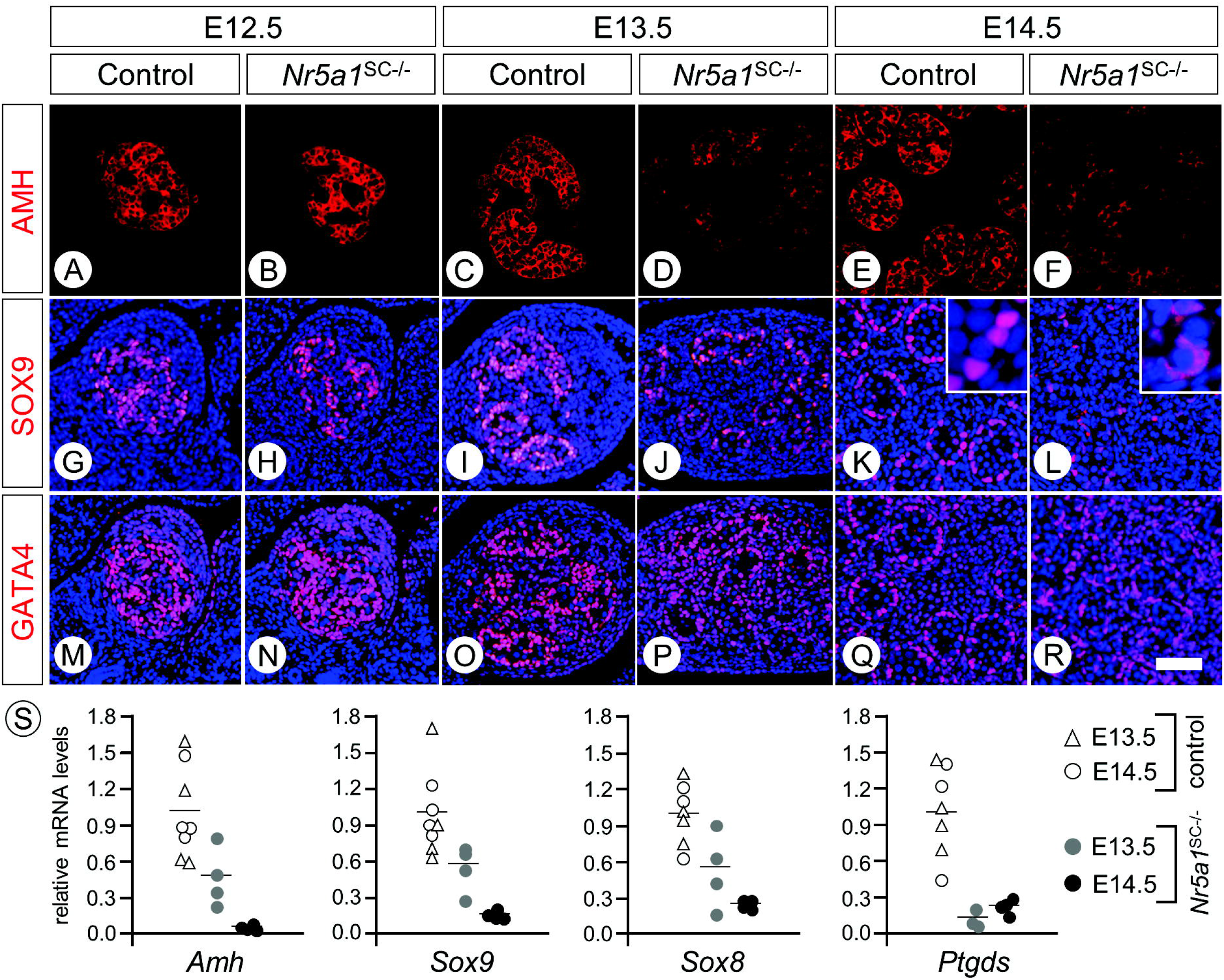
Ablation of *Nr5a1* in Sertoli cells impairs AMH and SOX9 expression. **(A-R)** Detection of AMH, SOX9 and GATA4 (red signals) on transverse histological sections of testes of control (A,C,E,G,I,K,M,O,Q) and *Nr5a1*^SC−/−^ mutant fetuses (B,D,F,H,J,L,N,P,R) at E12.5 (A,B,G,H,M,N), E13.5 (C,D,I,J,O,P) and E14.5 (E,F,K,L,Q,R). In (G-R) nuclei are counterstained with DAPI (blue signal). Insets (in K,L) are high magnifications showing nuclear localisation of SOX9 in control SC versus cytoplasmic localisation in mutant SC. Note that at each developmental stage, AMH, SOX9 and GATA4 IHC are performed on consecutive sections. Scale bar (in R): 50 μm. (**S**) RT-qPCR analyses comparing the expression levels of *Amh, Sox9, Sox8* and *Ptgds* mRNAs in whole testis RNA from control (n=4) and *Nr5a1*^SC−/−^ mutant (n=4) fetuses at E13.5 and E14.5. Each point represents the mean value of a technical triplicate, and the bars indicate the mean value.

In parallel, we analysed the expression of GATA4 and WT1, important for *Amh* expression and SC differentiation, respectively [19,20]. Both were detected at similar levels in SC nuclei of control and mutant testes, from E12.5 to E14.5 (**Fig.2M-R; Suppl.Fig.1**).

### *Nr5a1*-deficient SC die through a TRP53-independent mechanism

Thanks to the expression of the YFP-reporter we quantified the surface occupied by SC (green pixels) relative to the whole testis surface (**Fig.3A,D**). At E14.5, this ratio was about 30% lower in mutant than in control testes [28.3 ± 3.8 (n=5) versus 37.9 ± 2.5 (n=5), respectively; p<0.01]. This suggested that *Nr5a1*-deficient SC had either shut down expression of YFP or had disappeared from the testis. If YPF expression was shut down, the excised *Nr5a1* allele (L-) should be present in the testis at birth. If SC were lost, it should no longer be detected. PCR analysis of genomic DNA showed that the L– allele was not amplified in *Nr5a1*^SC−/−^ mutants (**Suppl.Fig.2A**), indicating that *Nr5a1*-deficient SC were lost from the mutant testis. In agreement with this possibility, numerous cells displaying features of SC (i.e., located at the periphery of the testis cords, amongst GATA4-expressing SC) were TUNEL-positive in E14.5 *Nr5a1*^SC−/−^ testis (arrowheads, **Fig.3E,F**). This indicates that NR5A1-deficient SC progressively died.

**Figure 3.**
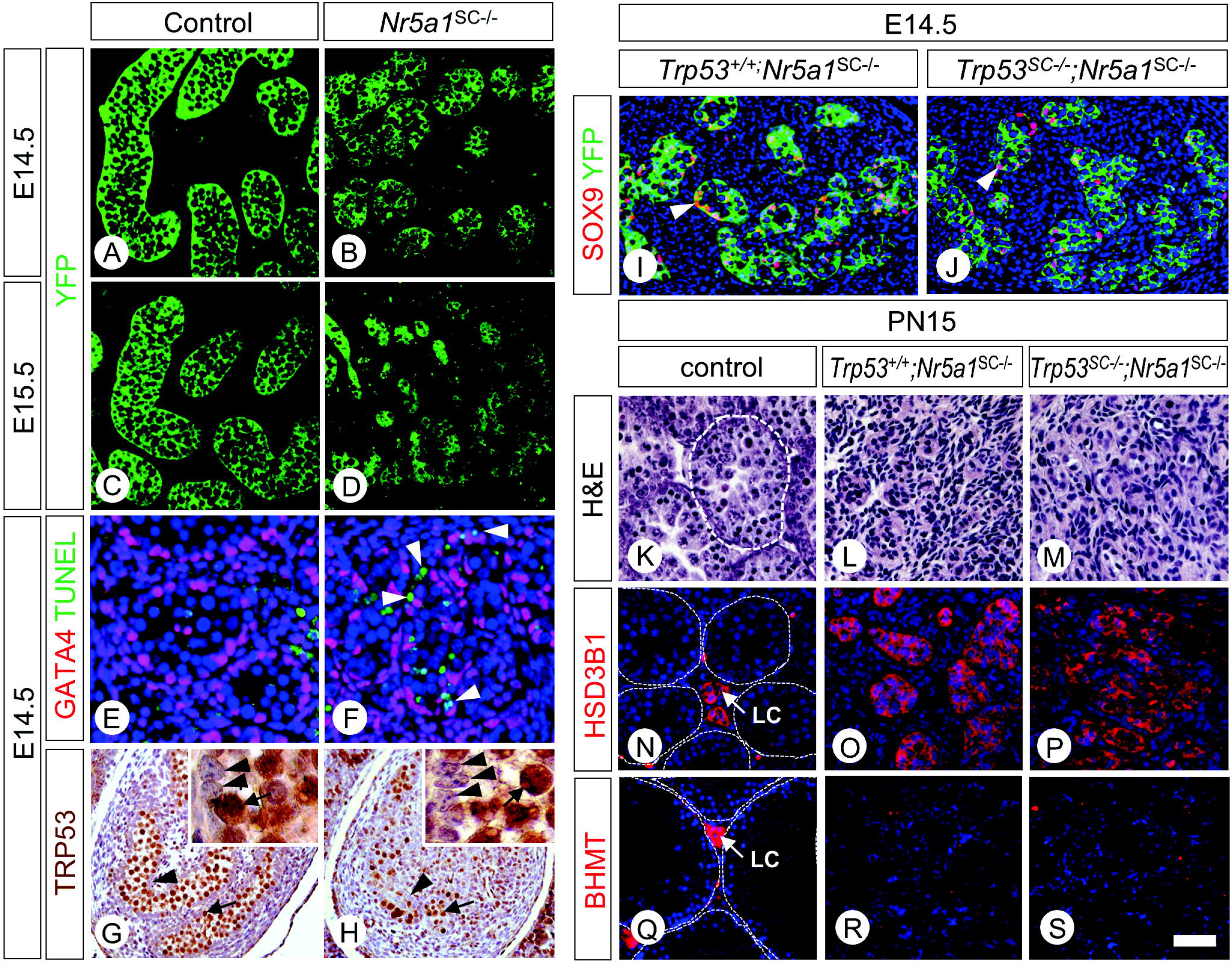
NR5A1-deficient Sertoli cells die even when they lack TRP53. **(A,D)** Detection of YFP (green signal) and TUNEL-positive cells (red signal) on histological sections from testis of control (A) and *Nr5a1*^SC−/−^ mutant fetuses (B) at E14.5 (A,B) and E15.5 (C,D). **(E,F)** Detection of GATA4 (red signal) and TUNEL-positive cells (green signal) on histological sections of control (E) and *Nr5a1*^SC−/−^ mutant testes (F) fetuses at E14.5. Nuclei are counterstained with DAPI (blue signal). Arrowheads point to TUNEL-positive SC in the mutant. **(G,H)** Detection of TRP53 (brown signal) on histological sections of control (G) and *Nr5a1*^SC−/−^ mutant (H) testes at E14.5. The insets show larger magnifications. Arrowheads and arrows point to SC and GC, respectively. **(I,J)** Detection of SOX9 (red signal) and YFP (green signal) by IHC on histological sections of *Nr5a1*^SC−/−^ (I) and *Nr5a1*^SC−/−^;*Trp53*^SC−/−^ (J) mutants at E14.5. Arrowheads point to SC. **(K-M)** Haematoxylin and eosin staining (H&E) on histological sections of testes from control (K), *Nr5a1*^SC−/−^ (L) and *Nr5a1*^SC−/−^;*Trp53*^SC−/−^ (M) mutants at PND15. **(N-P)** Detection of HSB3B1 (red signal) on transverse histological sections of testes from control (N), *Nr5a1*^SC−/−^ (O) and *Nr5a1*^SC−/−^;*Trp53*^SC−/−^ mutants (P) at PND15. **(Q-S)** Detection of HSB3B1 (red signal) on transverse histological sections of testes from control (Q), *Nr5a1*^SC−/−^ (R) and *Nr5a1*^SC−/−^;*Trp53*^SC−/−^ mutants (s) at PND60. The dotted white lines (in K,N,Q) delineate seminiferous tubules. Nuclei are counterstained with DAPI (blue signal). LC, Leydig cells. Scale bar (in S): 25 μm (A-D,I,J), 15 μm (E,F), 100 μm (G,H), 80 μm (K-P), 20 μm (Q-S).

Because TRP53 was not detected in fetal SC at E14.5 (**Fig.3G,H**), we wondered whether it was functionally involved in SC-death. We introduced conditional alleles of *Trp53* [21] in *Nr5a1*^SC−/−^ mutants bearing the YFP reporter transgene. Efficient ablation of *Trp53* was assessed by PCR on genomic DNA extracted form FACS-purified, YFP-positive, SC (**Suppl.Fig.2B**). E14.5 *Nr5a1*^SC−/−^ and *Nr5a1*^SC−/−^;*Trp53*^SC−/−^ mutants displayed similar reduced surfaces occupied by SC (**Fig.3I,J**, compare with 3A), out of which most of them had lost expression of SOX9. At post-natal day 15 (PND15), both *Nr5a1*^SC−/−^ and *Nr5a1*^SC−/−^;*Trp53*^SC−/−^ testes lacked SC-surrounded seminiferous tubules, but contained interstitial cells, out of which about half of them were HSD3B1-positive LC (**Fig.3K-P**). At adulthood, none of the interstitial cells expressed the adult LC-specific marker BHMT [22], indicating this cell-type was unable to emerge (**Fig.3Q-S**). Altogether, this indicates that NR5A1-deficient SC died, even though they lacked TRP53, yielding a postnatal testis made of FLC and other interstitial cells.

### *Nr5a1*^SC−/−^ adult males exhibit Müllerian duct retention

To analyse the outcome of the loss of SC and AMH production in *Nr5a1*^SC−/−^ mutants, the male phenotype was investigated at PND60. They all were sterile and had a shorter anogenital distance (AGD) index [0.30 ± 0.03 mm/g of body weight (n=16), versus 0.36 ± 0.02 mm/g (n=17) in controls; p<0.001] (**Fig.4A,C**). At autopsy (n = 8), they displayed normal derivatives of the Wolffian ducts (i.e., epididymis, vas deferent, seminal vesicle, ampullary glands) and the urogenital sinus (i.e., prostatic lobes) (**Fig.4E,F**). However, they also displayed Müllerian duct derivatives (**Fig.4F-H**), namely a vagina, an uterine body and bilateral, uterine horns, which were either incomplete and truncated (6 out of 8), or complete on one side (2 out of 8). Histological analyses revealed that the uterine body and vagina displayed a columnar epithelium (**Fig.4I)** and a stratified, squamous epithelium (**Fig.4J**), as anticipated for these female organs.

**Figure 4.**
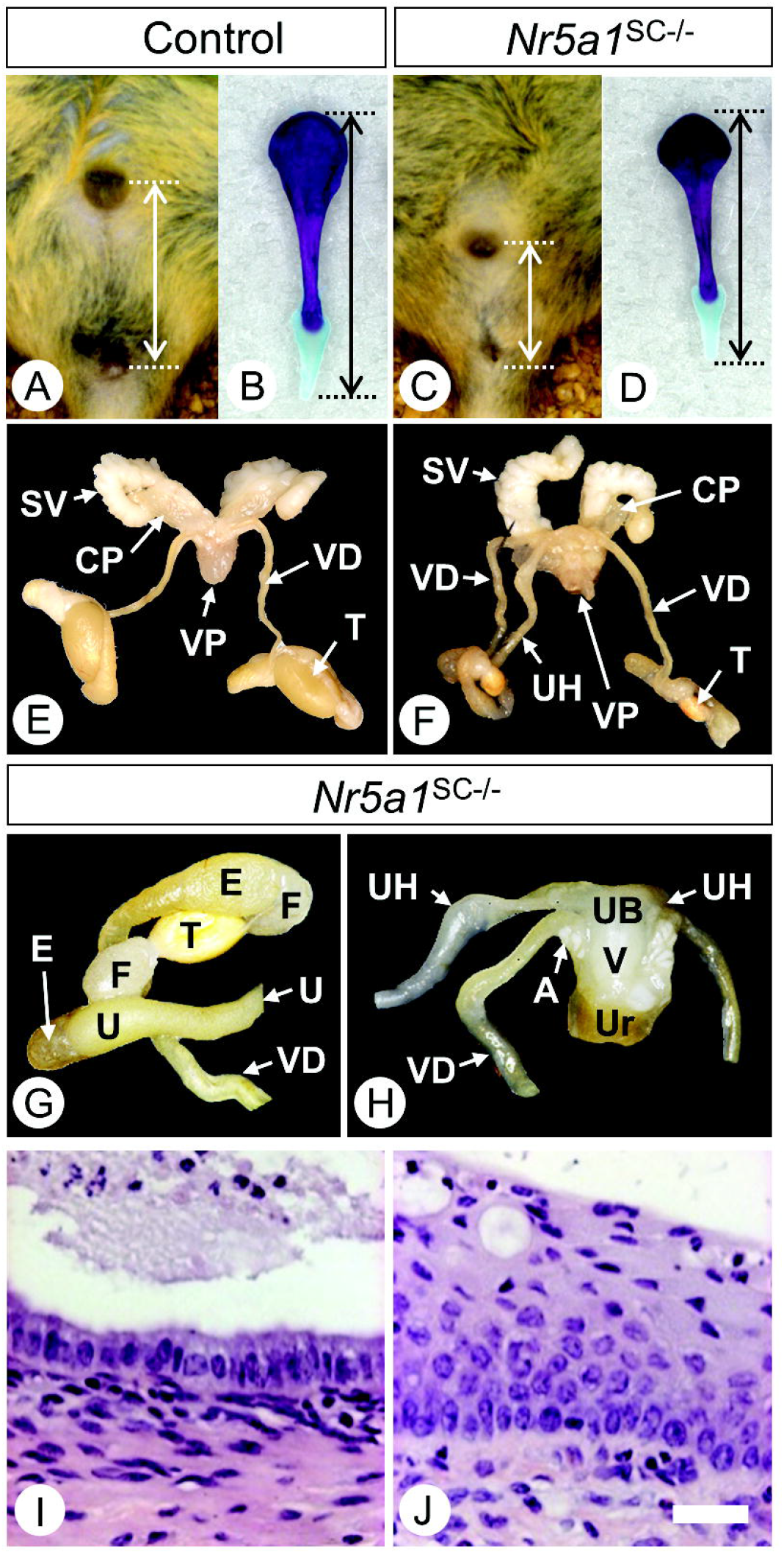
Abnormal external genitalia and Müllerian duct retention in *Nr5a1*^SC−/−^ males. **(A,C)** Anogenital distance in control and *Nr5a1*^SC−/−^ males at PND60. **(B,D)** Penis bones dissected from control and *Nr5a1*^SC−/−^ males at PND60 and stained with alizarin red and alcian blue. **(E-H)** Reproductive tracts of control (E) and *Nr5a1*^SC−/−^ (F-H) males at PN60. **(I-J)** Histological sections from the *Nr5a1*^SC−/−^ mutant sample shown in (H) and stained by H&E. The uterine surface consists of a simple columnar epithelium and the vaginal epithelium is typically stratified squamous. Legend: A, ampullary gland; CP, cranial prostate; E, epididymis; F, fat pad; SV, seminal vesicle; T, testis; UB, uterus body; UH, uterine horn; V, vagina; VD, vas deferens; VP, ventral prostate. Scale bar (in J): 10 μm (I,J).

Nonetheless, *Nr5a1*^SC−/−^ males displayed 40% lighter seminal vesicles [4.71 ± 1.66 mg/g of body weight (n=17) versus 7.43 ± 1.22 mg/g (n=16) in controls; p<0.05] and 10% shorter penis bones [6.7 ± 0.4 mm (n=17) versus 7.5 ± 0.3 mm (n=16) in controls; p<0.05] (**Fig.4B,D**). As AGD, seminal vesicle growth and penis bone length vary as a function of androgen exposure [23], we tested blood testosterone levels. They were comparable at birth [0.25 ± 0.17 ng/ml (n=10) in mutants, versus 0.22 ± 0.10 ng/ml serum (n=10) in controls; p=0.59], and at PND60 [0.38 ± 0.17 ng/ml (n=17) in controls, versus 0.43 ± 0.23 ng/ml serum (n=16) in controls; p=0.68], indicating normal testosterone production.

### Transcriptomic signatures of cells in control and *Nr5a1*-deficient gonads

We then performed single-cell RNA sequencing (scRNA-seq) experiments using dissociated cell suspensions obtained from 12 control and 16 *Nr5a1*^SC−/−^ whole testes at E14.5. After data processing and quality control, we assembled an atlas composed of 3,988 and 5,010 control and mutant testicular cells, respectively. On average, we detected ~13,860 unique molecular indices (UMIs) and ~3,556 genes in each individual cell. The 8,998 testicular cells were partitioned into 23 cell clusters (termed C1-C23) and projected into a two-dimensional space (**Fig.5A; Table.S1**). Known marker genes of distinct cell-types were used to identify each cluster (**Suppl.Fig.3**).

**Figure 5.**
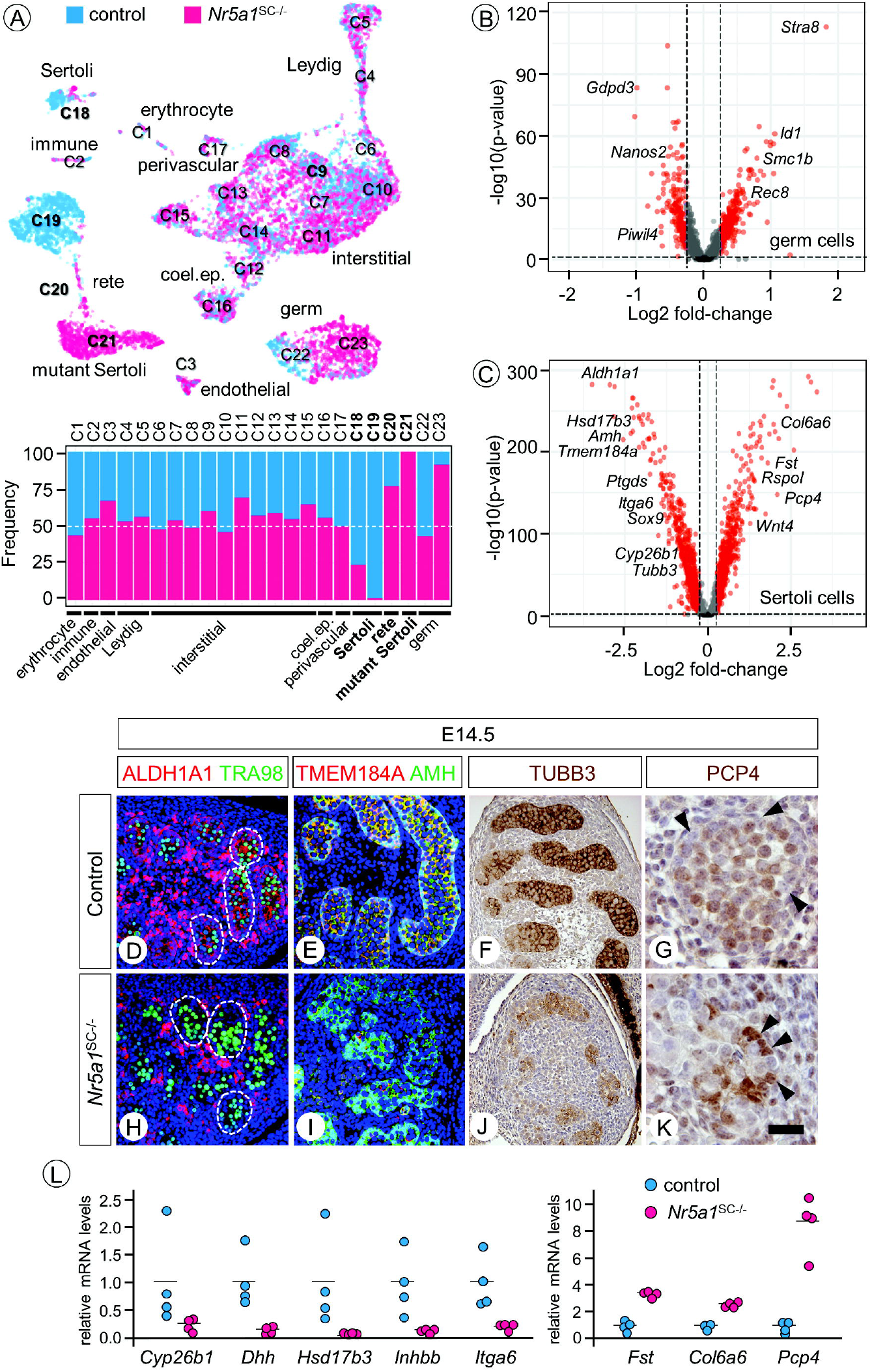
Transcriptomes of control and *Nr5a1*^SC−/−^ testicular cells at a single-cell resolution. **(A)** UMAP plot of single cell transcriptomes from control (blue) and *Nr5a1*^SC−/−^ (pink) testes at E14.5, partitioned into 23 cell clusters (named C1-C23) by using Seurat graph-based clustering. The proportion of control and mutant cells in each cell cluster is indicated below as coloured bars. Legend: Legend: coel. ep., coelomic epithelium. **(B,C)** Volcano plots of differential gene expression in GC (panel B, clusters C22 and C23) and SC (panel C, clusters C18, C19 and C21) from control and *Nr5a1*^SC−/−^ testes. Red dots correspond to genes deregulated more than 1.2-fold. **(D-K)** Detection of ALDH1A1, TMEM184A (red signals), TRA98, AMH, DDX4 (green signals) or TUBB3 and PCP4 (brown signals) by IHC on transverse histological sections of control (D-G) and *Nr5a1*^SC−/−^ mutant (H-K) testes at E14.5. In (D,E,H,I) nuclei are counterstained with DAPI (blue signal). The dotted lines (in D,I) delimit seminiferous cords. The arrowheads (in G,L) point to SC nuclei. Scale bar (in K): 25 μm (D,E,H,I), 100 μm (F,J), 15 μm (G,K). (**L**) RT-qPCR analyses comparing the expression levels of selected mRNAs as indicated in whole testis RNA from control (n=4) and *Nr5a1*^SC−/−^ mutant (n=4) fetuses at E14.5. Each point represents the mean value of a technical triplicate, and the bars indicate the mean value.

Only a few changes of gene expression were observed between control and *Nr5a1*^SC−/−^ testes for endothelial, immune and perivascular cells. In Leydig, interstitial, and coelomic epithelium cells the expression of 59, 49, and 47 genes was deregulated (**Table.S2**). Functional analysis revealed that down-regulated genes were related to ribosome in Leydig (14 genes, *p* value<4.8^e-15^) and interstitial cells (12 genes, *p* value<1.3^e-10^) (**Suppl.Fig.4; Table.S3**). In GC, the expression of 636 genes was significantly deregulated (**Table.S1**), amongst which *Stra8* and *Rec8* were up, while *Nanos2* and *Piwil4* were down (**Fig.5B**). Processes such as DNA recombination (33 genes, *p* value<2.0^e-10^), meiotic cell cycle (32 genes, *p* value<1.1^e-9^) and oocyte differentiation (8 genes, *p* value<5.6^e-3^) were identified amongst the upregulated genes (**Suppl.Fig.4; Table.S3**). In agreement with this finding we showed that GC were in the S-phase of the meiotic prophase at E14.5 and E15.5 in *Nr5a1*^SC−/−^ testes instead of becoming mitotically quiescent, as in the control situation. Accordingly, many GC of the *Nr5a1*^SC−/−^ testes expressed meiotic proteins, such as STRA8 or REC8, and died because of premature meiotic initiation (**Suppl.Fig.5**).

Three clusters of SC were identified (C18, C19 and C21), out of which C19 and C21 were mainly composed of control and NR5A1-deficient SC, respectively (**Fig.5A**). Almost all SC of C18 were mitotic (**Suppl.Fig.3**). In total, 1,671 genes were significantly deregulated in SC (**Suppl.Table.S1**), amongst which *Amh, Sox9* and *Ptgds* were down as anticipated (see above), while *Wnt4, Fst* and *Rspo1* were up (**Fig.5C**). We confirmed deregulated expression of some by IHC (**Fig.5D-K**) and that of others by RT-qPCR (**Fig.5L**), validating thereby the scRNA-seq experiment.

According to its transcriptomic signature, cluster C20 was assigned the *rete testis* identity (**Suppl.Fig.3)**. This structure is a set of ducts, through which spermatozoa are transported from the gonad via the efferent duct to the epididymis [24]. At E14.5, the *rete testis* is identified as epithelial cords between the mesonephric tubules and the testis cords (**Suppl.Fig.6A**). The *rete testis* cells express *Nr5a1, Pax8* and *Aldh1a3* [25]. As the *Plekha5*^Tg(AMH-cre)1Flor^ transgene is not functional in the *rete testis* [26], we anticipated that C20 could comprise cells in which NR5A1 expression was not lost. In agreement, we observed that cells in C20 organized as a continuum, which stemmed from C19 with *Nr5a1^+^/Yfp^−^/Pax8^−^/Aldh1a3*^−^ cells and extended with *Nr5a1^+^/Yfp^−^/Pax8^+^/Aldh1a3*^+^ cells (*rete*) originating from both control and *Nr5a1*^SC−/−^ testes. It ended at the junction with C21 with *rete testis* cells, originating exclusively from the *Nr5a1*^SC−/−^ testes, and in which cre was active (*Nr5a1^−^/Yfp^+^/Pax8^+^/Aldh1a3*^+^ cells) (**Suppl.Fig.6B**). Accordingly, IHC experiments confirmed that *rete testis* cells actually expressed NR5A1, but not YFP, control and *Nr5a1*^SC−/−^ testes (**Suppl.Fig.6C-J**).

### *Nr5a1*-deficient SC change their molecular identity

As indicated above, the NR5A1-deficient SC from cluster C21 gained expression of *Wnt4, Fst* and *Rspo1*, which characterize the fetal ovary somatic cells [4]. This raised the question as to whether SC had changed their identity. To test for this possibility, we decided to compare our data with a reference atlas of embryonic and fetal gonad development ranging from E10.5 to E16.5 [25]. After processing this atlas with the pipeline that we used for our data set, (**Suppl.Fig.7**), we next mapped our data set to this reference atlas (**Fig.6A**). Most of control and *Nr5a1*^SC−/−^ cell clusters were predicted to correspond to clusters with identical (or compatible) identities, as expected (**Fig.6B**). In addition, GC from control testes matched in the atlas with E13.5 male GC, while those from *Nr5a1*^SC−/−^ testes matched with E13.5 female (i.e., meiotic) GC (**Fig.6C**), as expected from IHC analyses (**Suppl.Fig.5**), and validating thereby the whole prediction. Interestingly, cluster C20 was predicted to match the best with *rete testis* cells, a feature fitting well with our analysis by IHC (see above, **Suppl.Fig.6**). Most importantly, cluster C21 was predicted to correspond to pGrC in the atlas. Further analysis distinguishing the developmental stages indicated that they actually matched with E16.5 pGrC (**Fig.6C**). The remaining cells matched with E13.5 SC. Thus, a fraction of NR5A1-deficient SC lost their identity, and acquired a “pGrC-like” identity. Accordingly, functional analysis of genes differentially expressed by NR5A1-deficient SC assigned upregulated genes to WNT signalling pathway (54 genes, *p* value<4.3e^−8^), known to drive pGrC differentiation in the fetal ovary [27] (**Suppl.Fig.4; Table.S3**). Nonetheless, the process of *trans*-differentiation was not fully achieved because IHC experiments failed to evidence expression of FOXL2 in the *Nr5a1*^SC−/−^ mutant testis at E15.5 (**Fig.6D,E**).

**Figure 6.**
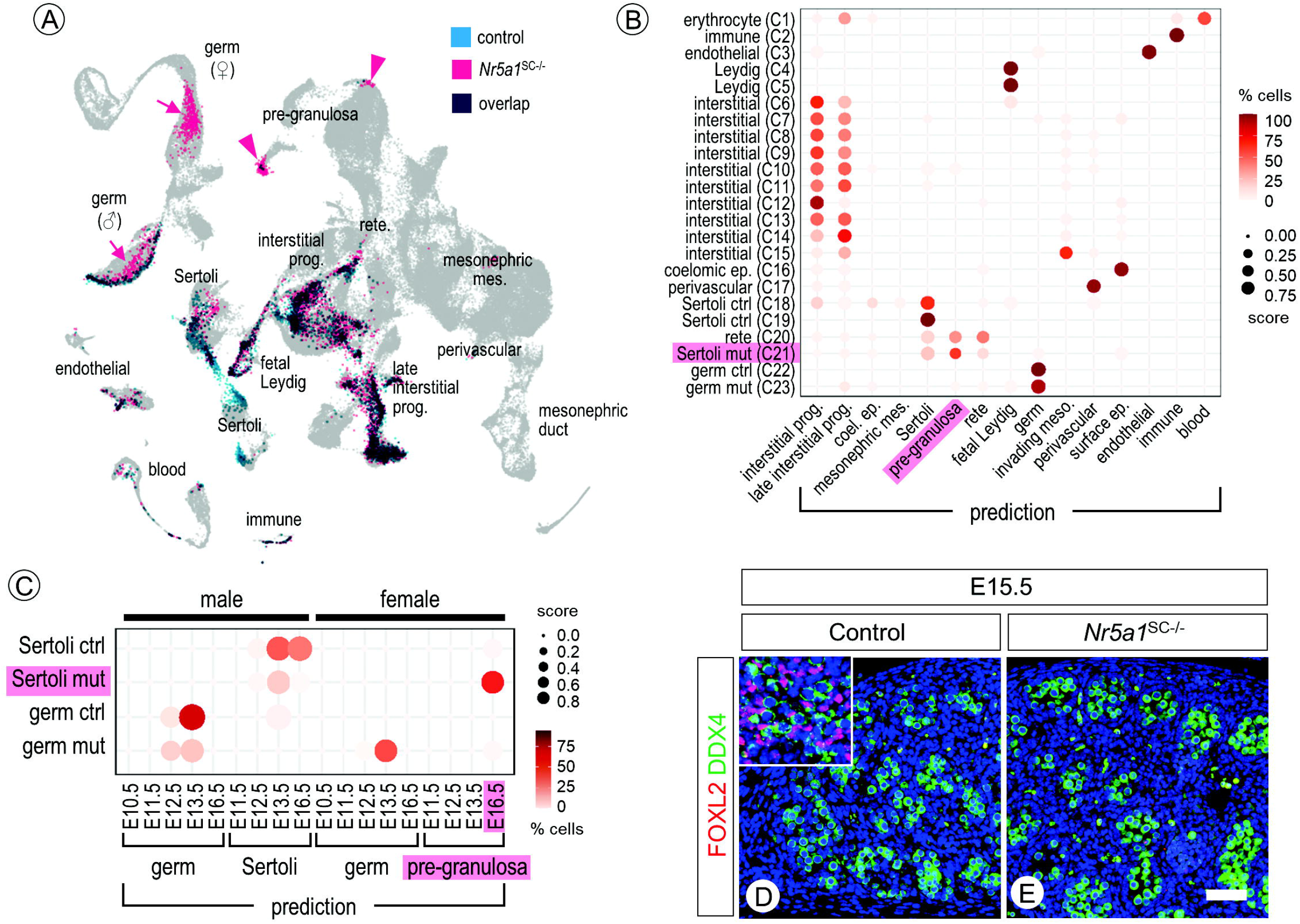
NR5A1-deficient Sertoli change their identity. **(A)** Predicted projection of control (blue) and *Nr5a1*^SC−/−^ (pink) cells on a reference single-cell transcriptomic atlas of gonad development. **(B,C)** Dot plots representing the predicted association of cell clusters from the control and *Nr5a1*^SC−/−^ single-cell dataset (y axis) to cell-types (panel B) and to sex and developmental stages (panel C) according to the atlas previously published [29] (x axis). The dot size represents the prediction score (ranging from 0.0 to 1.0) of a given cell cluster to be associated with a given cell-type based on the TransferData function implemented in Seurat. The color intensity (from white to dark red) indicates the percentage of cells of a given cell cluster that have been associated with a given cell-type. Legend: coel., coelomic; ctrl, control; ep., epithelium; mes., mesenchyme; meso., mesonephros; mut, mutant; prog., progenitor. **(D,E)** Detection of FOXL2 (red signal) and DDX4 (green signal) by IHC on transverse histological sections of control (D) and *Nr5a1*^SC−/−^ mutant (G) testes at E15.5. Nuclei are counterstained with DAPI (blue signal). The inset (in D) shows detection of FOXL2 in a E15.5 fetal ovary, used as a positive control for IHC. Scale bar (in E): 15 μm (D,E).

### Ablation of *Nr5a1* in SC induces disorganisation of the testis cords

On histological sections at E14.5, the majority of *Nr5a1*^SC−/−^ seminiferous cords displayed reduced diameters, as well as poorly defined and discontinuous contours (broken dotted line and arrows, **Fig.7E**). At E15.5, the cords had almost fully disappeared (broken dotted line, **Fig.7F**). Such irregularities were never observed in the control testes (**Fig.7A,B**).

**Figure 7.**
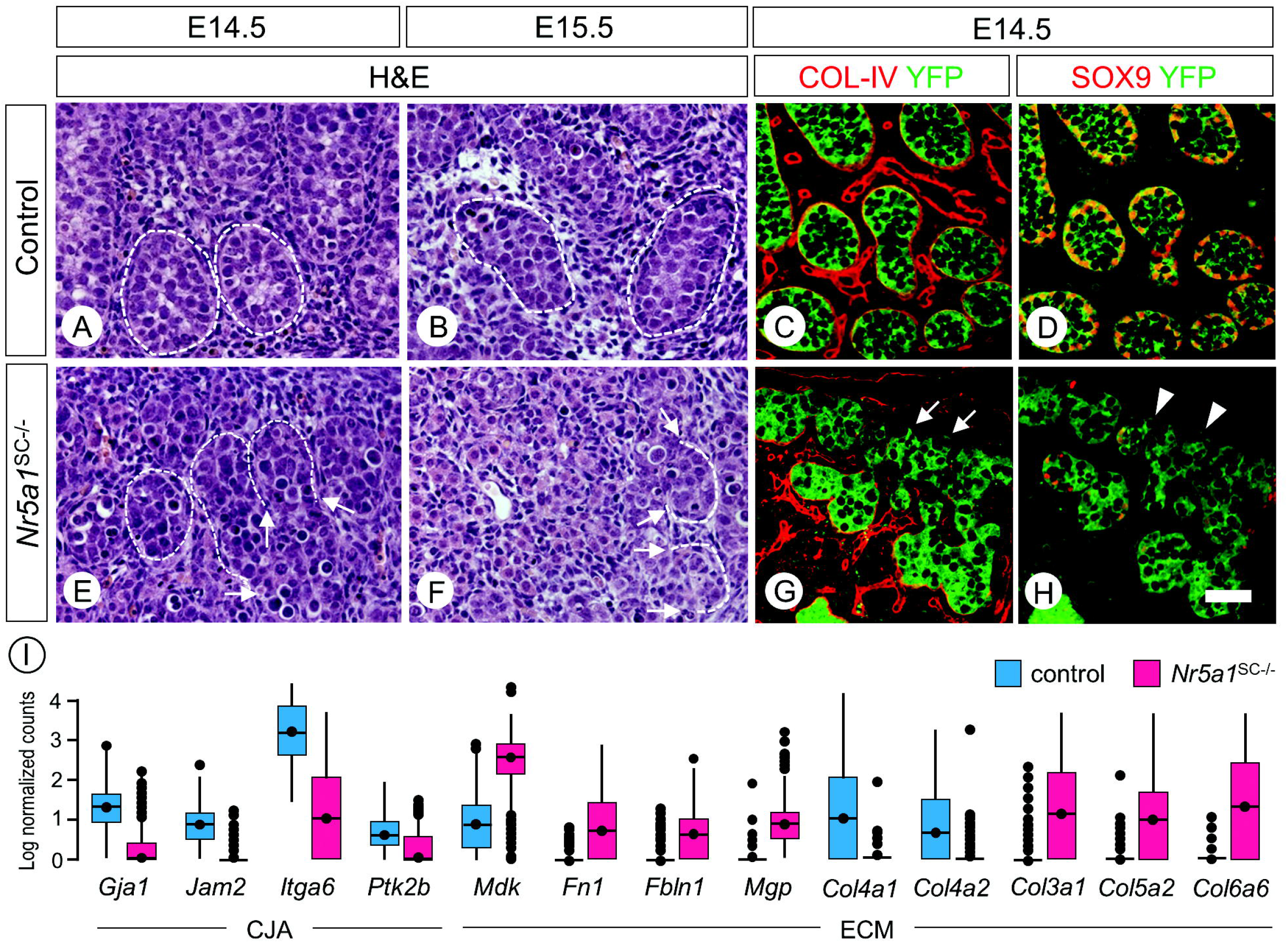
Loss of NR5A1 in Sertoli cells induces disorganisation of the testis cords. **(A,B,E,F)** Histological sections through E14.5 (A,E) and E15.5 (B,F) control (A,B) and *Nr5a1*^SC−/−^ mutant (E,F) fetuses stained by hematoxylin and eosin (H&E). In controls, the seminiferous cords are well defined (dotted lines). In mutants, they are poorly defined from E14.5 onwards (broken dotted lines with arrows). **(C,D,G,H)** Detection of COL-IV, SOX9 (red signals), and YFP (green signal) on transverse histological sections of the testis from control (C,D) and *Nr5a1*^SC−/−^ mutant fetuses (G,H) at E14.5. Note that (C,D) and (G,H) are consecutive sections. Arrows (in G) point to the loss of COL-IV at the periphery of seminiferous cords, where SOX9 is lost in SC (arrowheads in H). Scale bar (in D): 10 μm (A,B,E,F), 15 μm (C,D,G,H). **(I)** Tukey box plots illustrating medians, ranges and variabilities of log normalized expression of the indicated genes involved in either cell junction/adhesion (CJA) or ECM composition.

Importantly, functional analysis of the genes differentially expressed between control and NR5A1-deficient SC highlighted extracellular matrix (ECM; 74 genes, *p* value<2.6^e-16^), cell adhesion (118 genes, *p* value<5.9^e-9^), basement membrane (26 genes, *p* value<2.1^e-8^), as well as integrin binding (25 genes, *p* value<2.5^e-6^) (**Suppl.Fig.4; Table.S2**). Accordingly, the expression level of some genes involved in cell junction/adhesion (e.g., *Gja1, Jam2, Ptk2b*) and integrin signalling (e.g., *Itga6*) were down-regulated. As to genes for ECM components, some were down (e.g., *Col4a1, Col4a2*), while many others (e.g., *Col3a1, Col5a2, Col6a6, Fn1, Fbln1, Mdk, Mgp*) were up-regulated (**Fig.7I**). To verify this point, we analysed collagen type IV (COL-IV) expression by IHC. At E14.5, COL-IV was surrounding the whole periphery of all cords in control testes (yellow edging, **Fig.7C**). In *Nr5a1*^SC−/−^ testes, COL-IV was greatly reduced and even absent at the periphery of many cords (arrows, **Fig.7G**), in areas corresponding to regions where the expression of SOX9 was also lost (arrowheads, **Fig.7H**). As *Col4a1* and *Col4a2* are regulated by SOX9 [28], it is reasonable to propose that loss of SOX9 in NR5A1-deficient SC yielded loss of COL-IV and altered basement membrane, which, in turn, disorganized the cord structure. Searching further for ligand-receptor interactions in our data set, we identified a deregulated network related to integrin-dependent cell adhesion (**Suppl.Fig.8**). Together our findings indicate that NR5A1-deficient SC lost their appropriate cell-cell and cell-ECM interactions.

## DISCUSSION

Alteration of gene expression in *Nr5a1*^SC−/−^ testes could result from either loss of NR5A1 or death of SC. The decreased expression of *Sox9, Amh, Gja1*, and possibly *Cyp26b1* and *Dhh*, is likely directly related to the loss of NR5A1 in SC. Actually, NR5A1 initiates SC differentiation by directing *Sox9* expression [9]. Then, it maintains *Sox9* transcription and activates downstream genes by binding to *cis*-regulatory elements (e.g., *Amh, Gja1*) or synergizing through unknown mechanisms (e.g., *Cyp26a1, Dhh*) [16,29–32]. As to *Dmrt1, Sox8* and *Ptgds*, their decreased expression was also expected because SOX9 regulates their expression [17,31,33,34]. However, our finding that *Ptgds* is undetectable as early as E13.5, while SOX9 is still present, suggests that NR5A1 may regulate *Ptgds* expression directly (**Fig.8**).

**Figure 8.**
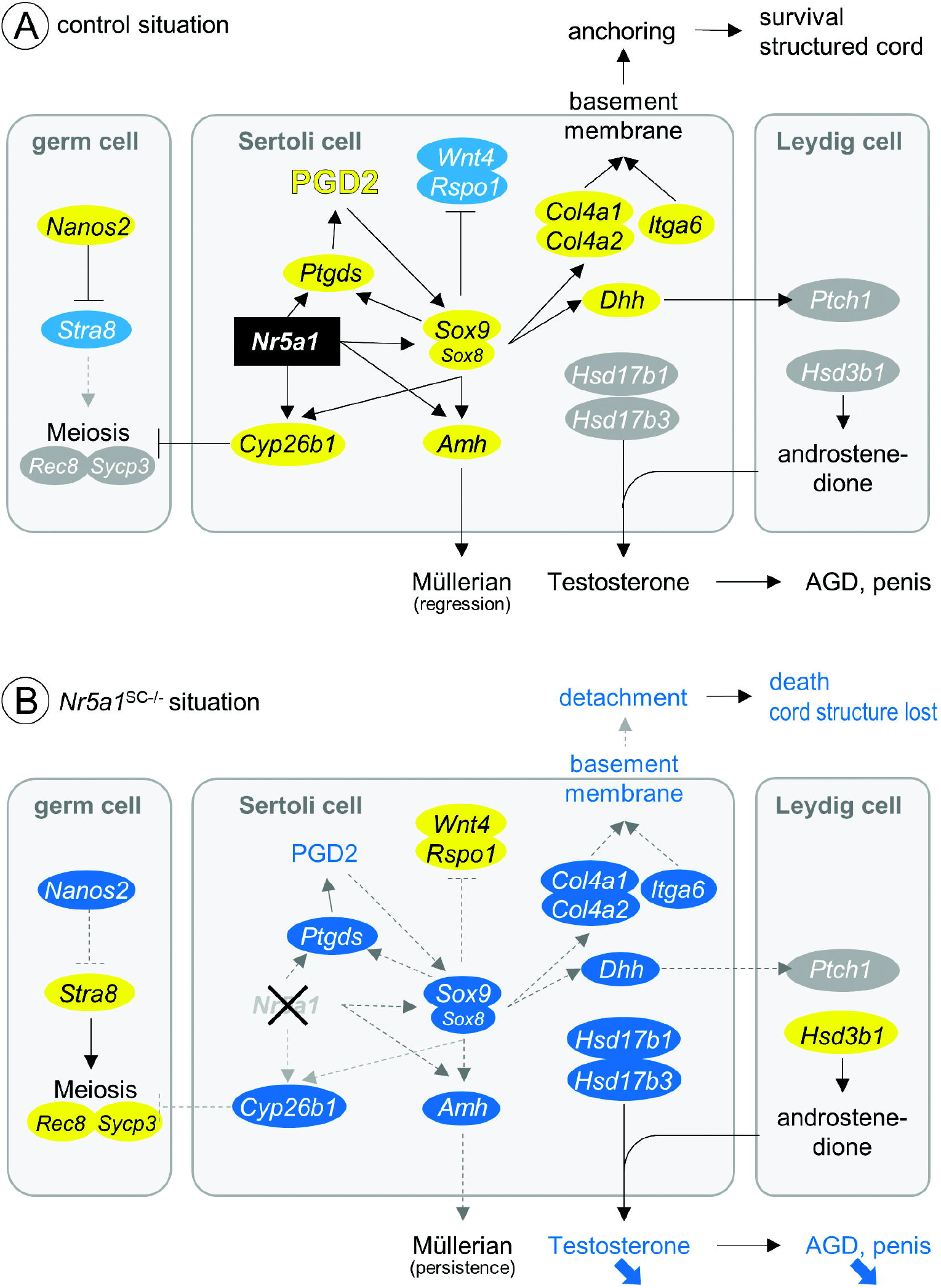
Summary of the alterations induced by the loss of NR5A1 in Sertoli cells after the sex determination period. **(A)** Control situation. **(B)** *Nr5a1*^SC−/−^ mutant situation. Blue and yellow ovals are up- and down-regulated genes, respectively. Legend: AGD, anogenital distance; PGD2, prostaglandin D2.

### NR5A1-deficient SC transdifferentiate into pre-granulosa-like cells

Based on genetic studies, SOX9 has emerged as the master regulator instructing SC differentiation during testis development, notably by repressing the WNT4/RSPO1-dependent pathways involved in pGrC differentiation [1–3]. However, human fibroblasts can be reprogramed into SC by forcing expression of NR5A1 and GATA4, but not SOX9 [35]. Here we show that SC lacking NR5A1 from E13.5 loose expression of genes defining or maintaining SC identity (e.g., *Sox9, Sox8, Ptgds, Dmrt1*), further illustrating the primary role of NR5A1 in instructing SC differentiation. In parallel, they gain expression of genes specifying pGrC differentiation (e.g., *Wnt4, Rspo1*) and their transcriptome projected to a gonad atlas clearly shows that they display an identity close to that of pGrC. Nonetheless, these “pGrC-like” cannot reach a functional state of differentiation since they keep FOXL2-negative. Another study has shown that SC lacking *Nr5a1* from E12.5 reach a functional, FOXL2-positive, granulosa cell identity, resulting in complete male-to-female sex reversal [36]. This indicates that SC devoid of NR5A1 from E12.5 can transdifferentiate into functional pGrC, while those loosing NR5A1 from E13.5 are no longer licensed to do so. We further show that these “pGrC-like” subsequently die through a *Trp53*-independent mechanism since concomitantly removing *Trp53* and *Nr5a1* does not prevent death. Quite interestingly, SC loosing *Nr5a1* later than E14.5 either die by MDM2/TRP53-dependent apoptosis or survive, since the resulting testis at adulthood still contains clearly defined seminiferous tubules [37]. Thus, the sooner NR5A1 is lost in SC, the highest their plasticity remains, suggesting that NR5A1 progressively locks the SC identity over time. In addition, SC die by distinct mechanisms, depending on the stage at which NR5A1 is lost.

### Sertoli cells lacking NR5A1 from E13.5 die by anoikis

It is accepted that cell adhesion/junction molecules and ECM components play important roles in maintaining the integrity of seminiferous cords [38–40]. Because many genes deregulated in NR5A1-deficient SC are coding for such molecules, including collagen-coding genes, testis cord disruption is likely linked to breakdown of SC adhesion to their environment. Interestingly enough, some of the upregulated genes coding for ECM components such as *Col18a1, Fn1, Nid2* and *Dcn* (**Table.S2**) are characteristics of pGrC [41].

Cells sense their location and maintain their adhesion through specific interactions with the ECM. This plays a major role in regulating various processes, including cell-survival. This function is mediated by integrins, which are the cell-membrane receptors interacting with components of the ECM. They recruit nonreceptor tyrosine focal adhesion kinases (FAK), that are activated in response to adhesion [42]. Therefore, disruption or loss of integrin-ECM adhesion impairs cell-survival, and often causes detachment-induced cell death, called anoikis [43,44]. We found that expression of *Itga6*, an integrin subunit crucial for fetal testis organization [45], and *Ptk2b*, a member of the FAK family expressed in SC in a SOX9-dependent manner [46] is reduced in NR5A1-deficient SC. Thus, it is conceivable that these changes may cause SC to lose their anchorage to the modified ECM (see above) and initiate death of detaching SC by anoikis, altering thereby integrity of the testis cord epithelium (**Fig.8**). It is worth noting that anoikis still occurs in the absence of TRP53 [47], which is fully consistent with our finding that NR5A1-deficient SC die independently of TRP53.

### NR5A1-deficient SC and differentiation of the other testicular cell-types

As to the GC fate, the data show that most of them become meiotic and die in *Nr5a1*^SC−/−^ testes. The fact that expression of *Cyp26b1* is lost in NR5A1-deficient SC is sufficient to explain why GC entered meiosis (**Fig.8**). Actually, CYP26B1 is required in SC to prevent meiotic initiation and to maintain GC in an undifferentiated state [48].

As to FLC, their specification and/or development rely on SC, notably thanks to DHH- and PDGF-signalling pathways [49,50]. Our finding that FLC are present in *Nr5a1*^SC−/−^ testes, despite decreased expression of *Dhh* and *Pdgfa* in SC, reveals that these pathways are no longer required beyond E13.5 to allow proper emergence or survival of FLC, as observed when SC are ablated after E15.5 [51,52]. Furthermore, these FLC appear functional since testicular descent is occurring in *Nr5a1*^SC−/−^ mutants, suggesting normal INSL3 production [53].

As to peritubular myoid cell (PTMC), it is admitted that SC are required during the fetal period to support their differentiation, notably by producing DHH and ECM. Genetic ablation of SC from E14.5 or *Dhh*-null mutations actually induce seminiferous cords defects closely resembling those observed in *Nr5a1*^SC−/−^ testes [54–56]. Thus, the decreased expression of *Dhh* in NR5A1-deficient SC and their death questions the fate of PTMC. As a matter of fact, we could not distinguish PTMC from the interstitial or coelomic epithelium cell clusters. This is not really surprising as PTMC, sparse at E14.5, become obvious only from E16.5 [57], i.e., 2 days later than the stage at which we performed analysis. It is nonetheless worth noting that expression of *Ptch1, Tagln, Acta2* and *Tpm1*, which are generally considered as hallmarks of PTMC [58,59], were significantly reduced in interstitial or coelomic epithelium cluster cells (**Table.S2**). It is therefore possible that PTMC, included in these clusters, are altered, or even lost, following ablation of *Nr5a1* in SC. Interestingly, both PTMC and “dormant” steroidogenic progenitors (SP) at the origin of adult LC stem from a single population of *Wnt5a+* SP [60]. In this context, it is conceivable that the absence of adult LC in PND60 testis is related to an alteration of PTMC during fetal development. Further investigations are required to trace the fate of these *Wnt5a+* SP in *Nr5a1*^SC−/−^ testis.

### Death of the NR5A1-deficient SC affects genital tract development

Müllerian duct regression in males proceeds rostro-caudally, and is dependent on the action of AMH, which begins as early as E13.5 [61,62]. As *Amh* expression in *Nr5a1*^SC−/−^ mutants is lost only from E14.5, AMH production starts normally and then gradually decreases to zero after E14.5. It is therefore not surprising that the rostral part of the Müllerian ducts regresses normally, making their derivatives (oviducts, anterior portions of the uterine horns absent in the mutants. On the other hand, it is reasonable to propose that the caudal part of the Mullerian ducts persists beyond E14.5, and is at the origin of the posterior portions of the uterine horns, body of the uterus and the vagina of adult *Nr5a1*^SC−/−^ mutants (**Fig.8**).

In addition, *Nr5a1*^SC−/−^ mutants display normal Wolffian duct-derived genitalia (epididymis, vas deferens, seminal vesicles) [63]. These develop under the influence of testosterone, the synthesis of which starts as early as E13.5 in mice [64]. Their presence indicates therefore that *Nr5a1*^SC−/−^ mutants were exposed to testosterone during fetal development. This is surprising since SC, converting FLC-produced androstenedione into testosterone thanks to expression of *Hsd17b1* and *Hsd17b3* [65], are progressively lost in *Nr5a1*^SC−/−^ mutants. Thus, another enzyme necessarily compensates for the loss of *Hsd17b1* and *Hsd17b3*, as considered elsewhere [66]. In this regard, the HSD17B12 is a good candidate since it can convert androstenedione to testosterone [67]. However, the fact that FLC are not able to produce testosterone from androstenedione [65] means one has to consider HSD17B12 allows testosterone synthesis in a cell-type other than FLC. This production is nevertheless insufficient since AGD and penile bone length, both of which are highly sensitive to testosterone levels between E14.5 and E17.5 in mice [68], are reduced in *Nr5a1*^SC−/−^ mutants (**Fig.8**).

Our study shows that SC lacking NR5A1 from E13.5 transiently transdifferentiate to acquire a pre-granulosa cell-like identity, profoundly modify the landscape of the adhesion molecules and ECM they express and die by a TRP53-independent mechanism related to anoikis. This yields a disorganization of the testis cords, making GC to prematurely enter meiosis instead of becoming quiescent. Interestingly, the maintenance of fetal LC is not altered in the *Nr5a1*^SC−/−^ mutant, but the emergence of adult LC is clearly compromised. Further studies are now required to understand the reason for the absence of adult LC in the postnatal testis.

## MATERIALS AND METHODS

### Mice

They were on a mixed C57BL/6 (50%)/129/SvPass (50%) genetic background. The *Nr5a1* conditional mutant mouse line was established at the Institut Clinique de la Souris (*i*CS, Illkirch, France), in the context of the French National Infrastructure for Mouse Phenogenomics PHENOMIN (http://www.phenomin.fr). To construct the targeting vector a 1.9 kb-long DNA fragment encompassing exon 7 (ENSMUSE00000 693512) was amplified by PCR using 129/SvPass genomic DNA and cloned into an *i*CS proprietary vector containing a *lox*P site, as well as a *lox*P- and *FRT*-flanked neomycin resistance cassette (step1 plasmid). Then, 3 kb- and 3.7 kb-long fragments corresponding to 5’ and 3’ homology arms were amplified by PCR and introduced into step1 plasmid to generate the targeting construct. This linearized construct was electroporated into 129/SvPass mouse embryonic stem (ES) cells. After selection, targeted clones were identified by PCR using external primers and confirmed by Southern blots (5’ and 3’ digests) hybridized with neomycin, 5’ and 3’ external probes. One positive ES clone was injected into C57BL/6J blastocysts. To remove the selection cassette from the *Nr5a1* locus, chimeric males were crossed with *Gt(ROSA)26Sor^tm1(FLP1)Dym^* females [69]. Germline transmission was obtained, and a further breeding step was needed to segregate animals bearing the *Nr5a1* L2 allele from animals bearing the transgene. To inactivate *Nr5a1* in SC, female mice bearing *Plekha5*^Tg(AMH-cre)1Flor^ [14] and *Gt(ROSA)26Sor*^tm1(EYFP)Cos^ [15] transgenes, and heterozygous for the L2 allele of *Nr5a1* were mated with males heterozygous or homozygous for L2 alleles of *Nr5a1*. The resulting *Plekha5*^Tg(AMH-cre)1Flor^;*Nr5a1*^+/+^; *Gt(ROSA)26Sor*^tm1(EYFP)Cos^ and *Plekha5*^Tg(AMH-cre)1Flor^;*Nr5a1*^L2/L2^; *Gt(ROSA)26Sor*^tm1(EYFP)Cos^ males are referred to as control and *Nr5a1*^SC-/-^ mutant fetuses, respectively. To test for the role of TRP53, the L2 allele of *Trp53* gene [21] was further introduced in the mice described above.

Noon of the day of a vaginal plug was taken as 0.5 day embryonic development (E0.5). All fetuses were collected by caesarean section (no randomization, no blind experiments). Yolk sacs or tail biopsies were taken for DNA extraction. Primers 5’-GTCAAGCGCCCCATGAATGC-3’ and 5’-TTAGCCCTCCGATGAGGCTG-3’ were first used to amplify *Sry* gene (230 bp-long fragment) for male sex determination. Then, primers 5’-TGAGCCCTGGCACATCCCTCC-3’ and 5’-CCTCTGCCCTGCAGGCTTC TG-3’ were used to detect *Plekha5*^Tg(AMH-cre)1Flor^ transgene (273 bp-long amplicon), and primers 5’-AAGGGAGCTGCAGTGGAGTA-3’ and 5’-GCCAGAGGCCACTTGTGTAG-3’ to detect *Gt(ROSA)26Sor^tm1(EYFP)Cos^* reporter (520 bp-long amplicon). Primers 5’-CTGTCTCCTGTCTTCTACTACCCTG-3’ and 5’-AGCCATTTCAACAGTGCCCC TTCC-3’ were used to amplify wild-type (+, 290 bp-long) and L2 (400 bp-long) alleles of *Nr5a1*, while primers 5’-GTGGCACATGCATTAGTCCACTTGG-3’ and 5’-AGCCATTTCAACAGTGCCCCTTCC-3’ were used to amplify the excised, null, L– (243 bp-long) allele. Primers 5’-CACAAAAACAGGTTAAACCCAG-3’ and 5’-AGCACATAGGAGGCAGAGAC-3’ were used to amplify the wild-type (288 bp-long) and L2 (370 bp-long) alleles of *Trp53*. Primers 5’-CACAAAAACAGGTTAAACCCAG-3’ and 5’-GAAGACAGAAAAGGGGAG GG-3’ were used to amplify the excised, null, L– (612 bp-long) allele of *Trp53*. The PCR conditions were 30 cycles with denaturation at 95°C for 30 seconds, annealing at 61 °C for 30 seconds and elongation at 72°C for 30 seconds. The amplicons were resolved on 1.5% (w/v) agarose gels, stained by ethidium bromide and visualized under UV light, using standard protocols.

### Blood sample collection and testosterone measurement

After anesthesia with a lethal dose of Xylasin and Ketamine as described above, the blood of 9-11-week-old adult mice was collected into heparinized Microvette tubes (Sarstedt, Nümbrecht, Germany) by intra-cardiac sampling. The tubes were then centrifuged at 5000 g for 5 minutes and the resulting plasma samples were frozen until further use. Testosterone concentrations were determined by ELISA using a commercially available kit AR E-8000 (LDN, Labor Diagnostika Nord, Nordhorn, Germany). Statistical significance was further assessed by using two-tail Student’s *t*-tests.

### Morphology, histology and immunohistochemistry

Adult mice (PND60) were anesthetized by intraperitoneal injection of a lethal anesthetic mixture made of Xylasin (3 mg/ml) and Ketamine (20 mg/ml), and tissues were immediately fixed by intracardiac perfusion of 4% (w/v) paraformaldehyde (PFA) dissolved in phosphate buffered saline (PBS). Following collection, E12.5-E15.5 fetuses and tissues were fixed for 16 hours in 4% (w/v) PFA at 4°C or in Bouin’s fluid at 20°C. After removal of the fixative, samples were rinsed in PBS and placed in 70% (v/v) ethanol for long-term storage, external morphology evaluation and organ weight measurement. They were next embedded in paraffin and 5 μm-thick sections were made. For histology, sections were stained with hematoxylin and eosin (H&E).

For lHC, antigens were retrieved for 1 hour at 95°C either in 10 mM sodium citrate buffer at pH 6.0 or in Tris-EDTA at pH 9.0 [10 mM Tris Base, 1 mM EDTA, 0.05% (v/v) Tween 20]. Sections were rinsed in PBS, then incubated with appropriate dilutions of the primary antibodies (**Table.1**) in PBS containing 0.1% (v/v) Tween 20 (PBST) for 16 hours at 4°C in a humidified chamber. After rinsing in PBST (3 times for 3 minutes each), detection of the bound primary antibodies was achieved for 45 minutes at 20°C in a humidified chamber using Cy3-conjugated or Alexa Fluor 488-conjugated antibodies. Nuclei were counterstained with 4’,6-diamidino-2-phenylindole (DAPI) diluted at 10 μg/ml in the mounting medium (Vectashield; Vector Laboratories, Newark, CA, USA). ImmPRESS® Polymer Detection Kits MP-7500 and MP-7405 were used according to the manufacturer’s protocol (Vector Laboratories).

**Table 1.**
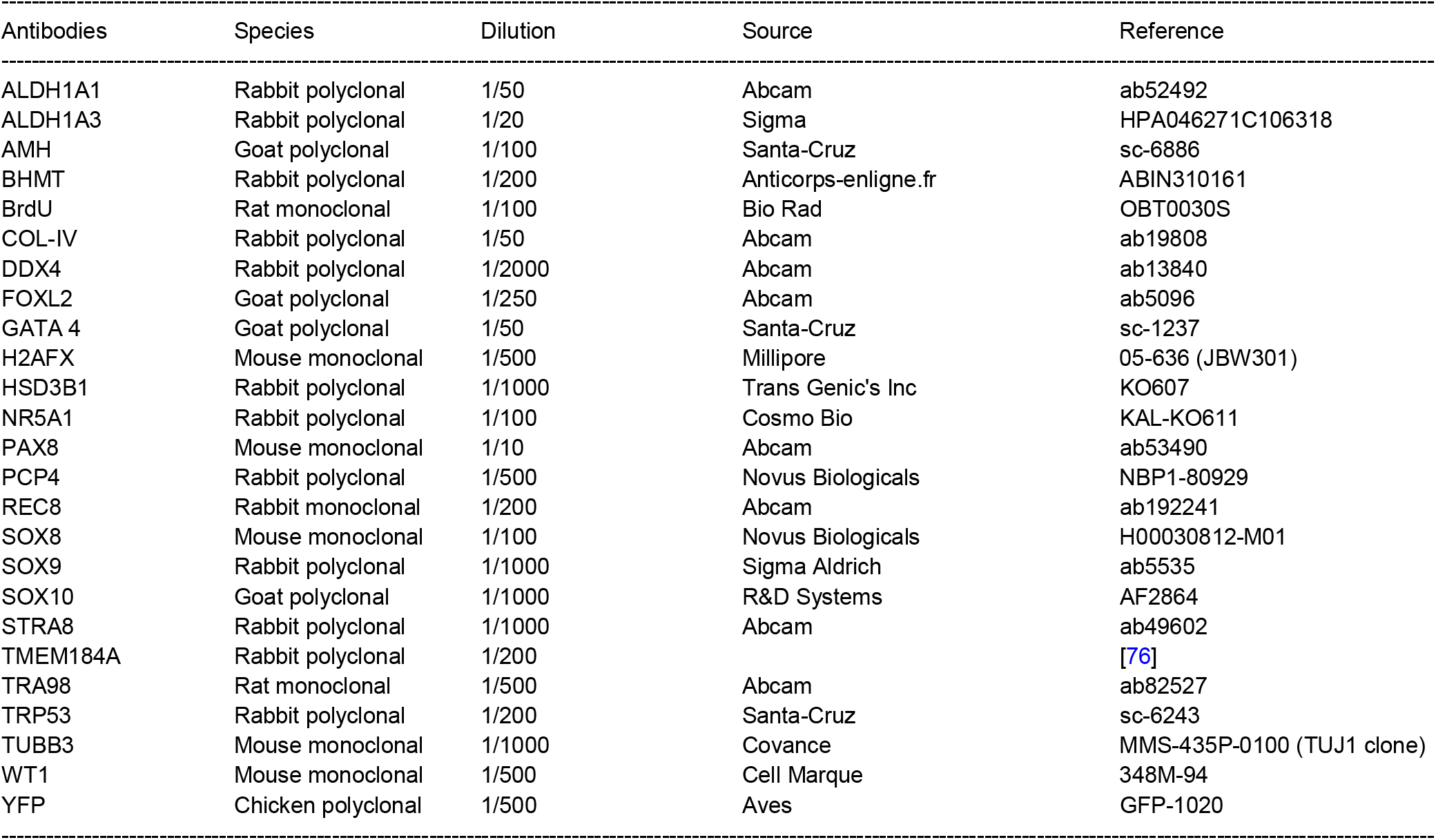
Primary antibodies used for immunohistochemistry experiments.

The surface area occupied by YFP-positive cells was measured using a macro command designed for Fiji software (Imaging Center of IGBMC). Data were expressed as percentage of YFP-positive surface areas relative to the entire testis section surface areas. At least four samples were analyzed per genotype. Statistical analysis was done by a two-tail Student *t*-test, assuming equal variances after arcsine transformation of the percentages.

### BrdU incorporation and TUNEL assays

BrdU (Sigma-Aldrich, Saint-Quentin-Fallavier, France) dissolved at 5 mg/ml in PBS was injected intraperitoneally to pregnant females at 50 mg/kg of body weight. Two hours later, fetuses were collected (E13.5-E15.5), fixed and embedded as described above. BrdU incorporation was detected on 5 μm-thick sections by using an anti-BrdU mouse monoclonal antibody (diluted 1:100) and indirect IHC as described above. At least three samples per stage and per genotype were analyzed. Data were expressed as percentages of BrdU-positive cells related to the number of DDX4-positive cells. Statistical analysis was done by a two-tail Student *t*-test, assuming equal variances after arcsine transformation of the percentages.

TUNEL-positive cells were detected on sections from PFA-fixed samples using the In Situ Cell Death Detection Kit, Fluorescein, according to the manufacturer’s instructions (Roche, Mannheim, Germany). At least three samples were analyzed per genotype. Data were expressed as the ratio between the number of TUNEL-positive cells quantified on entire sections and the surface areas of the testis sections (μm^2^). Statistical significance was assessed by using two tail Student’s *t*-tests.

### Real-time RT-qPCR analyses of RNA extracted from whole gonad

Fetal testes were dissected, isolated from mesonephros, snap frozen in liquid nitrogen and stored at −80°C until use. Whole testis total RNA was extracted using RNeasy Mini Kit (Qiagen, Les Ulis, France). RT-qPCR was performed on 5 ng RNA aliquots using Luna® Universal One-Step RT-qPCR Kit, according to the manufacturer’s instructions (New England Biolabs, Evry, France). The primers are listed in **Table.2**. Triplicates of at least four samples were used for each genotype, at each stage. The relative transcript levels were determined using the ΔΔCt method, and normalized to *Hprt* whose expression is not affected by ablation of *Nr5a1*.

**Table 2.**
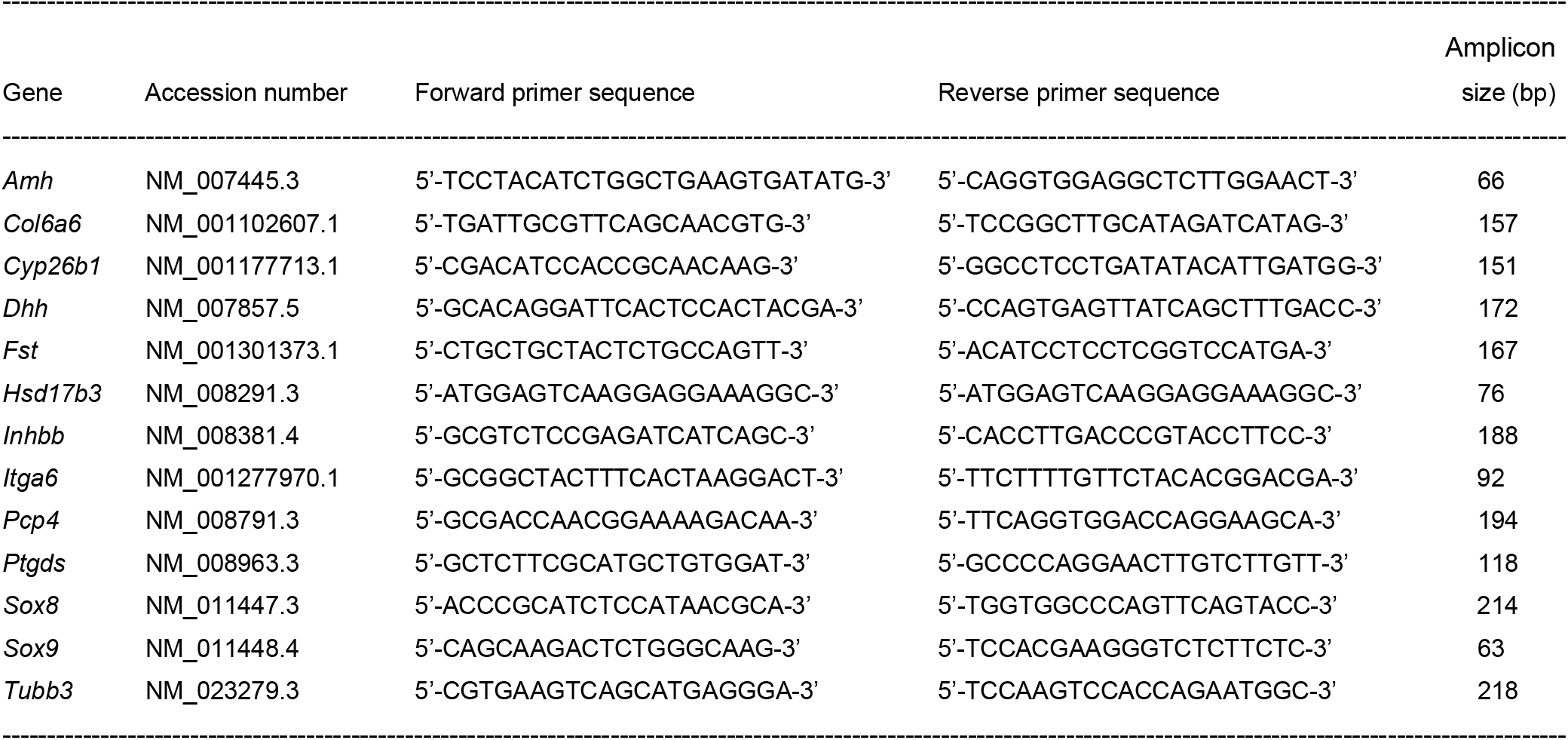
Primers used for quantification of mRNA levels by real-time RT-qPCR.

### Purification of YFP-positive Sertoli cells

To dissociate cells, the testes of controls and mutants additionally bearing the *Gt(ROSA)26Sor*^tm1(EYFP)Cos^ transgene were incubated for 10 minutes at 37°C in 350 μl of trypsin/EDTA 0.05% (w/v), phenol red solution (Gibco Invitrogen, Auckland, New-Zealand), filtered through a 40 μm cell strainer to generate single cell suspensions, centrifuged at 3000g and suspended in 300 μl PBS as described [70]. The YFP-positive and -negative cells were sorted separately by FACS using an Aria® II flow cytometer (BD Biosciences, Le Pont de Claix, France). Sorted cell suspensions were then lysed for overnight at 55°C in a proteinase K-containing buffer, DNA was extracted and genotyped by PCR using standard protocols and primers as indicated above.

### Single cell RNA sequencing and data processing

Gonads from E13.5 and E14.5 control and mutant fetuses were dissected out in PBS, sexed by their appearance under the microscope and cell suspensions were prepared as described above. Cell number and viability were determined by a Trypan Blue exclusion assay on a Neubauer Chamber. Samples consisting of > 90% viable cells were processed on the Chromium Controller from 10X Genomics (Leiden, The Netherlands). Ten thousand cells were loaded per well to yield approximately 5000 to 6000 captured cells into nanoliter-scale Gel Beads-in-Emulsion (GEMs). Single cell 3’ mRNA sequencing libraries were generated according to Chromium Single Cell 3’ Reagent Kits User Guide (v3 Chemistry) from 10X Genomics. Briefly, GEMs were generated by combining barcoded gel beads, a reverse transcription master mix containing cells, and partitioning oil onto Chromium Chip B. Following full length cDNA synthesis and barcoding from poly-adenylated mRNA, GEM were broken and pooled before cDNA amplification by PCR using 11 cycles. After enzymatic fragmentation and size selection, sequencing libraries were constructed by adding Illumina P5 and P7 primers (Illumina, Evry, France), as well as sample index via end repair, A tailing, adaptor ligation and PCR with 10 cycles. Library quantifications and quality controls were determined using Bioanalyzer 2100 (Agilent Technologies, Santa Clara, CA, USA). Libraries were then sequenced on Illumina HiSeq 4000™ as 100 bases paired-end reads, using standard protocols.

Sequencing data were processed using 10X Genomics software (https://support.10xgenomics.com/single-cell-gene-expression/software/overview/welcome). Fastq files were processed with CellRanger (v6.0) on a chimeric genome composed of mm10 *Mus musculus* assembly and 753 bp-long YFP sequence (European Nucleotide Archive accession number AGM20711) [71]. The resulting count matrices were aggregated with the Read10X function implemented in Seurat (v4.0.1) [72]. Cells with less than 200 detected genes and genes detected in less than 10 cells were removed. Doublets were filtered out independently in each individual matrix by using the DoubletFinder R package (v.2.0.2) [73]. Subsequently, the Single-Cell Analysis Toolkit for Gene Expression Data in R (scater v1.10.1) was used to remove outlier cells by using several cell features including the proportions of reads mapping mitochondrial and ribosomal genes, the number of genes and UMIs per cell [74]. Data were normalized using the NormalizeData and the SCT function implemented into Seurat by regressing out the unwanted variation due to cell cycle and to the proportion of mitochondrial genes. The top-3000 most varying genes were used to perform a principal component analysis with the RunPCA function implemented in Seurat. Cells were then clustered by using Seurat graph-based clustering (FindNeighbors and FindClusters functions) on the top-30 principal components, with default parameters. Finally, we used the Uniform Manifold Approximation and Projection (UMAP) method (RunUMAP function) to project cells in a 2D space. Cell clusters were annotated using a set of known marker genes. The FindAllMarkers function implemented in Seurat was used to identify significantly differentially expressed genes (DEGs) between cell clusters. Gene ontology (GO) and pathway enrichment analysis was conducted for each gene expression cluster using the AMEN suite of tools [75] with an BH-adjusted p value of ≤ 0.05. The current dataset was mapped on top of a reference atlas of gonadal development [25] using FindTransferAnchors and MapQuery functions with default settings. To address this issue, the reference atlas was first processed by using the same pipeline.

## ACKNOWLEDGEMENTS

The *Nr5a1* mouse mutant line was established at the Institut Clinique de la Souris (PHENOMIN *i*CS) in the Genetic Engineering and Model Validation Department with funds from the ANR (see below). We thank Pr Anne-Marie LEFRANCOIS-MARTINEZ for discussions and advices. We also thank Dr Christelle Thibault-CARPENTIER from the Genomeast platform for her input (http://genomeast.igbmc.fr/). This research was funded by Agence Nationale pour la Recherche Grants (see below). A CC-BY-4.0 public copyright license has been applied by the authors to the present document and will be applied to all subsequent versions up to the Author Accepted Manuscript arising from this submission, in accordance with the grant’s open access conditions.

## AUTHOR INFORMATION

These authors contributed equally: Frédéric Chalmel, Norbert B. Ghyselinck.

### Contributions

SSC, MM and NBG conceptualized and designed the study. SSC and DC performed all the experiments. BF and MK provided technical and material support. EG contributed to image analysis. SSC, VA, MC, MJ and FC performed the scRNA-seq experiment and analysed data. CM and SN provided datasets for the mouse gonad atlas. SSC, NV, MM, FC and NBG analysed all experiments, managed the overall study, wrote the manuscript, and supervised its preparation and submission. NV, MM, FC and NBG acquired funding for this research.

### Corresponding authors

Correspondence to <u>Norbert B. Ghyselinck</u>.

## ETHICS DECLARATIONS

### Ethics statement

Mice were housed in a licensed animal facility (agreement #C6721837). All experiments were approved by the local ethical committee (Com’Eth, accreditation APAFIS#18323-2018113015272439_v3), and were supervised by N.B.G., M.M. and N.V., who are qualified in compliance with the European Community guidelines for laboratory animal care and use (2010/63/UE).

## DATA AVAILABILITY

The data sets described in this paper are available at Gene Expression Omnibus (GEO) database under the accession number GSE219271.

## SUPPLEMENTARY INFORMATION

Supplementary information is available at *Cell Death and Differentiation’s* website.

