## Supplementary information for "Loss of NR5A1 in Sertoli cells after sex determination changes their cellular identity and induces their death by anoikis"

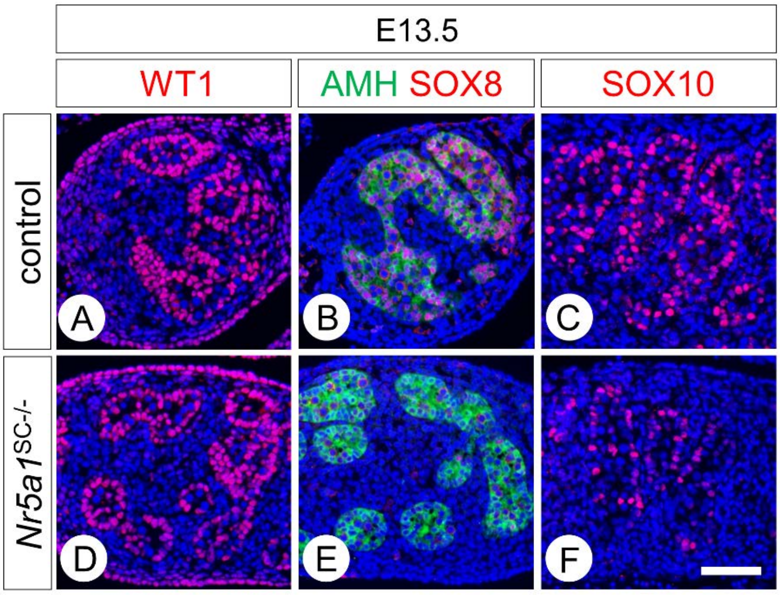


**Supplementary Figure 1. Ablation of *Nr5a1* in Sertoli cells (SC) impairs SOX8 and SOX10 expression. (A-F)** Detection of WT1, SOX8, SOX10 (red signals) and AMH (green signals) on transverse histological sections of the testis of a control (A-C) and a *Nr5a1*$^{SC–/–}$ mutant fetus. (D-F) at E13.5. Nuclei are counterstained with DAPI (blue signal). Scale bar (in F): 50 µm (A-F).

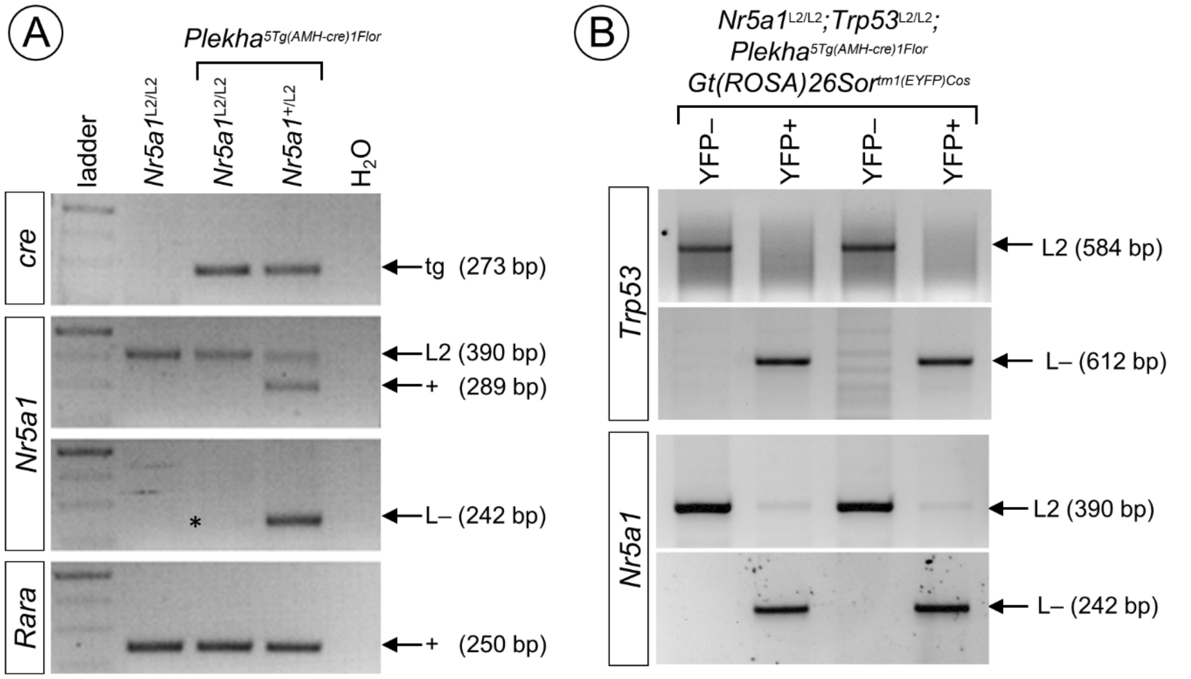


**Supplementary Figure 2. NR5A1-deficient Sertoli cells (SC) die even without TRP53 (A)** PCR analysis of genomic DNA extracted from control (*Nr5a1*$^{L2/L2}$), mutant (*Plekha5*$^{Tg(AMH\text{-}cre)1Flor}$; *Nr5a1*$^{L2/L2}$) and heterozygote (*Plekha5*$^{Tg(AMH\text{-}cre)1Flor}$; *Nr5a1*$^{+/L2}$) testes of newborns. Upper panel shows genotyping of *Cre* transgene; middle panel shows genotyping of *Nr5a1* alleles; lower panel shows genotyping of *Rara* locus, attesting for an equivalent loading of DNA in each lane. The sizes of the expected fragments are indicated on the right: tg, Cre transgene; L2 and +, *lox*P-flanked and wild-type alleles, L–, excised, null allele. Note that the mutant testis contains only traces of the *Nr5a1* L– allele (asterisk), indicating that SC bearing the excised allele are no longer present at birth. **(B)** PCR analysis of genomic DNA extracted from FACS-purified YFP-positive (YFP+) and negative (YFP) cells contained in *Nr5a1*$^{SC–/–}$ ;*Trp53*$^{SC–/–}$ mutants fetuses at E14.5. Left panel shows genotyping of *Nr5a1* alleles; right panel shows genotyping of *Trp53* alleles. The YFP-negative cells (i.e., all cells except SC) contain only the unexcised (L2) alleles, while the YFP-positive cells (i.e., SC) contain only the cre-recombined, null (L–) alleles. Note however that traces of the *Nr5a1* L2 alleles are detected in YFP-positive cells.

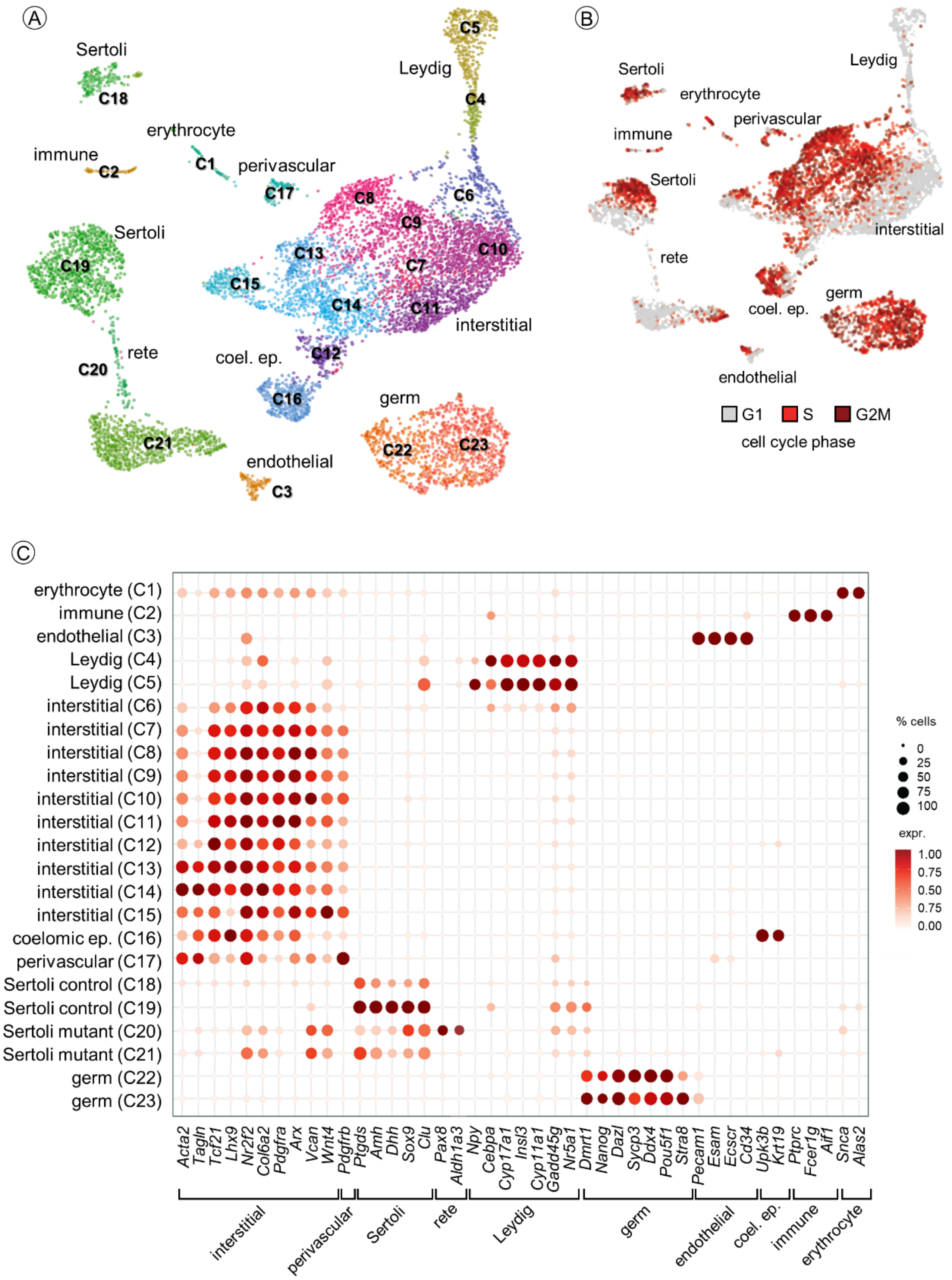


**Supplementary Figure 3. Identification of cell clusters generated from the single-cell transcriptomes. (A-B)** UMAP projection of the 8,998 cells coloured by cell clusters (panel A) or by phases of the cell-cycle, as indicated (panel B). Associated cell annotation is indicated close to the corresponding cell clusters (named C1-23). **(C)** Dot plot with the expression of selected markers (x axis) for each cell cluster (y axis). The dot size represents the percentage of cells expressing a given gene within a given cell cluster. The colour intensity (from light to dark red) indicates the average expression (log normalized counts) of a given gene within a given cell cluster.

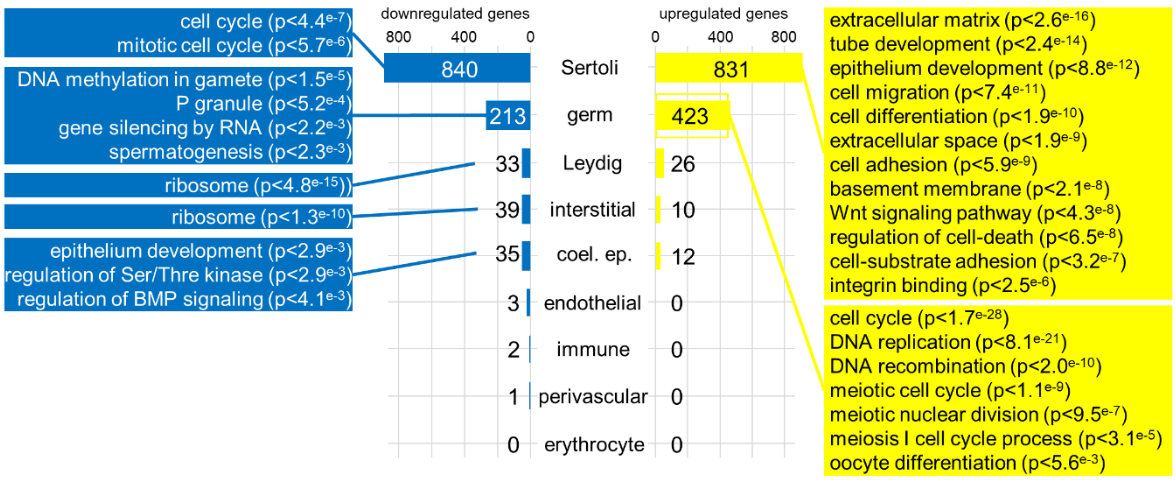


**Supplementary Figure 4. Number of differentially expressed genes and associated GO terms**. Bar plot representing the number of significantly downregulated (left side) and upregulated (right side) genes between control and *Nr5a1*$^{SC-/-}$ cells (x axis), in each cell-type (y axis). Enriched GO terms and their associated *p* values are given in each cell-type. Downregulated or upregulated genes and their associated GO terms are blue and yellow coloured, respectively.

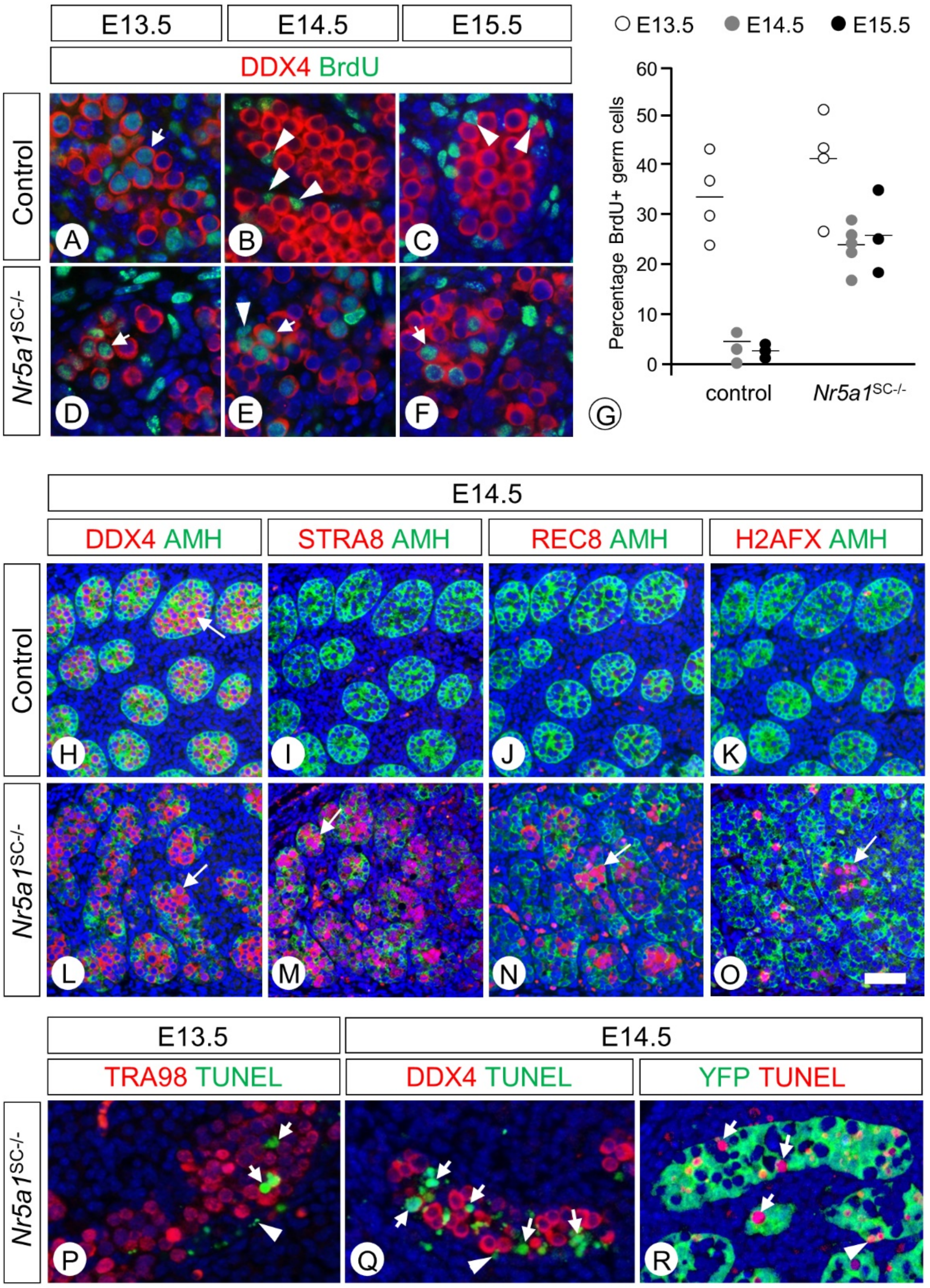


**Supplementary Figure 5. Germ cells (GC) initiate meiosis and die in *Nr5a1*$^{SC–/–}$ mutants, even in the absence of *Trp53*. (A-F)** Detection of DDX4 (red cytoplasmic signal) and BrdU (green nuclear signal) by IHC on histological sections of control (A-C) and *Nr5a1*$^{SC–/–}$ mutant (D-F) testes at E13.5 (A,D), E14.5 (B,E) and E15.5 (C,F). Arrows and arrowheads point to BrdU-positive GC and SC, respectively. **(G)** Dot plots showing the percentage of BrdU-positive GC in control (n=3 to 4) and mutant (n=3 to 4) testes, as a function of the developmental stages. **(H-O)** Detection of DDX4, STRA8, REC8, H2AFX (red signals) and AMH (green cytoplasmic signal) by IHC on histological sections of control (H-K) and *Nr5a1*$^{SC-/-}$ mutant (L-O) testes at E14.5. Arrows point to GC. The meiotic proteins STRA8, REC8, H2AFAX are detected in the mutant but not in the control testes. **(P,Q)** Detection of TUNEL-positive cells (green signal) and TRA98- or DDX4-positive GC (red signals) on histological sections from E13.5 (P) and E14.5 (Q) *Nr5a1*$^{SC-/-}$ mutant testes. **(R)** Detection of TUNEL-positive cells (red signal) and YFP-positive SC (green signal) on histological sections from an E14.5 mutant testis. The total number of TUNEL-positive cells was significantly larger in *Nr5a1*$^{SC-/-}$ testes than in controls [79 ± 9 cell/mm$^2$ (n=6) versus 27 ± 11 cell/mm$^2$ (n=3), respectively; $p<0.05$]. Scale bar (in O): 10 µm (A-F,P,R), 50 µm (H-O).

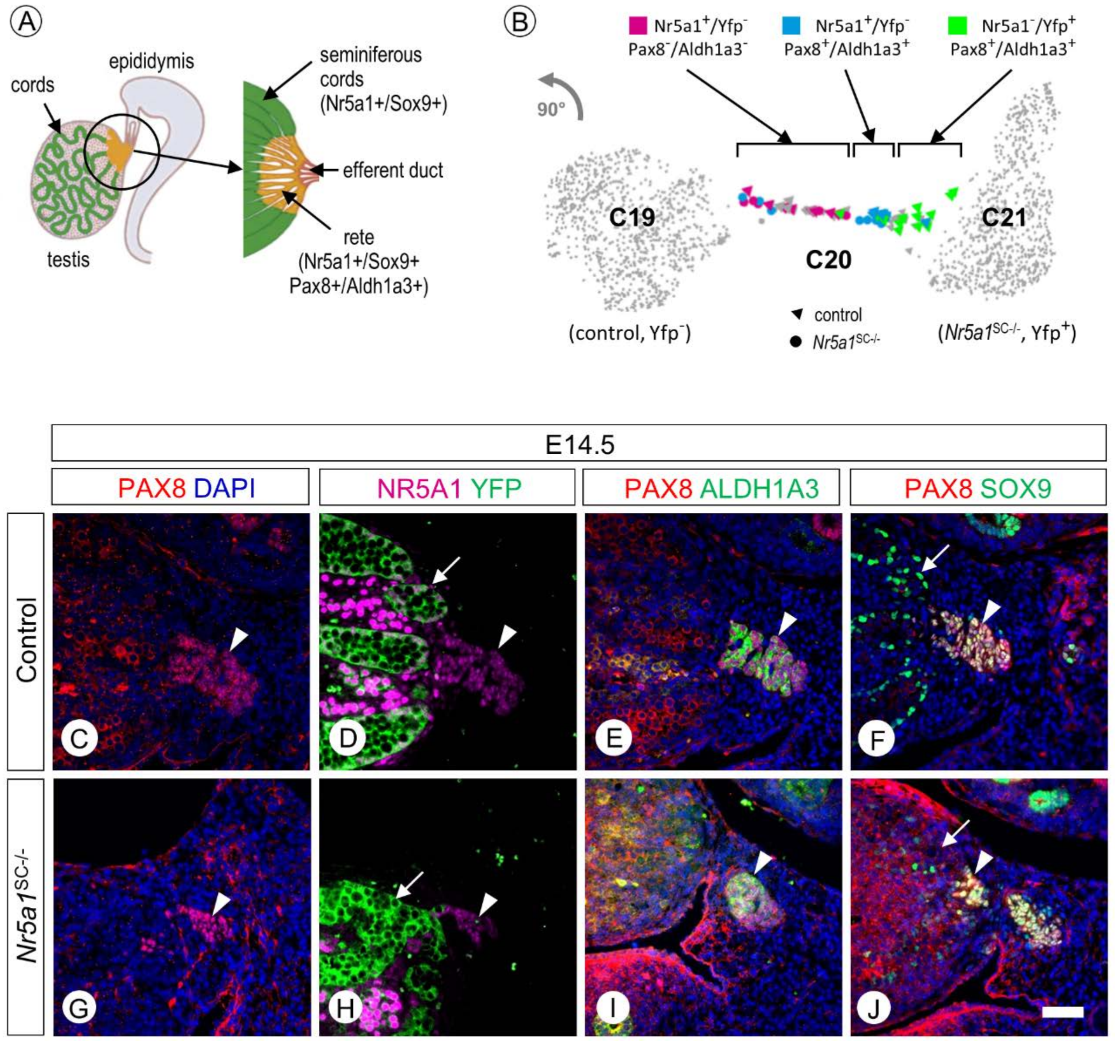


**Supplementary Figure 6. Cluster C20 corresponds to *rete* testis cells, in which gene excision by cre recombinase is not operational**. **(A)** Diagram illustrating the location and organization of *rete* testis in the mouse. Seminiferous cords are in green, while *rete* testis is in brown. **(B)** Magnification of UMAP plot (rotated by 90° with respect to Fig. 4) for clusters C19, C20 and C21. Cells belonging to C19 (from control testes, YFP-negative cells) and to C21 (from *Nr5a1*$^{SC–/–}$ testes, YFP-positive) are in grey. Cells belonging to C20 and expressing distinct combination between *Nr5a1*, *Yfp*, *Pax8* and *Aldh1a3* are depicted by a colour code: pink represents cells expressing *Nr5a1* but not *Yfp*, *Pax8* or *Aldh1a3*; blue stands for cells expressing *Nr5a1*, *Pax8* and/or *Aldh1a3*, but not *Yfp*; green represents cells expressing *Yfp*, *Pax8* and/or *Aldh1a3*, but not *Nr5a1*. Triangles and circles represent cells from control and *Nr5a1*$^{SC–/–}$ testes, respectively. **(C-J)** Detection of PAX8 (red signal) and YFP, ALDH1A3 or SOX9 (green signals) by IHC on histological sections of control (A-F) and *Nr5a1*$^{SC–/–}$ mutant (G-J) testes at E14.5. Arrows and arrowheads point to SC and *rete* testis cells, respectively. The *rete* testis cells (arrowheads) are NR5A1-positve, YFP-negative and SOX9-positive in both control (D,F) and mutant testes (H,J). In contrast, SC (arrows) are NR5A1-negaive, YFP-positive and SXO9-negative in the mutant (H,J), but not in the control testes (D,F). Scale bar (in J): 50 µm (C-J).

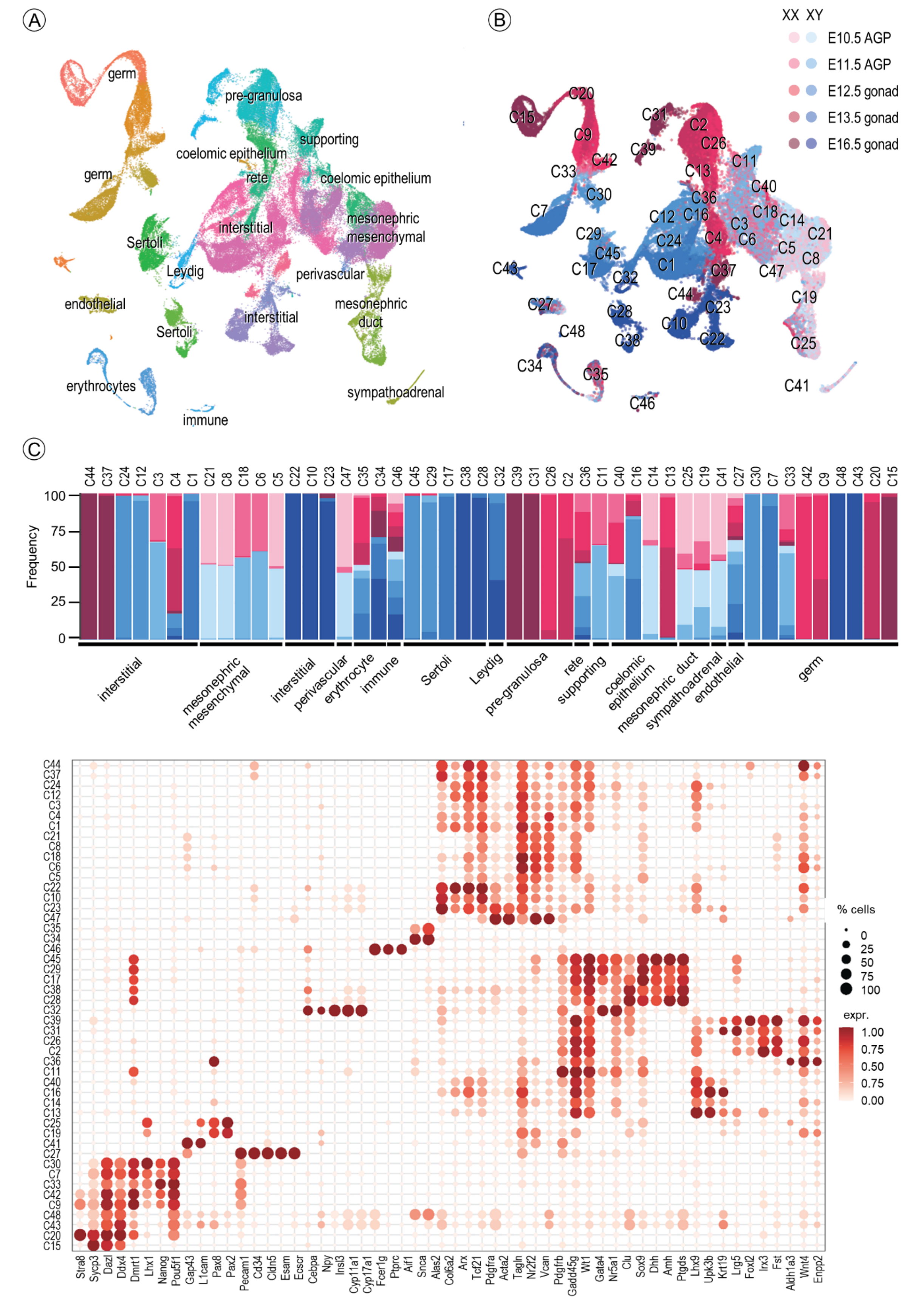


**Supplementary Figure 7. Identification of cell clusters generated from the atlas. (A-B)** UMAP projection of the 94,705 cells from the single-cell transcriptomic atlas of gonad development recently published [25] coloured by cell clusters (in panel A) or by developmental stage, from E10.5 to E16.5 as indicated (in panel B). Cell clusters (C1-C48) and their associated cell annotation are indicated**. (C)** Stacked bar plot showing the proportion of the developmental stages (from light at E10.5 to dark colour at E16.5) amongst the different cell clusters with male (XY, blue) and female (XX, pink) cells. (D) Dot plot with the expression of selected markers (x axis) for each cell cluster (y axis). The dot size represents the percentage of cells expressing a given marker within a given cell cluster. The colour intensity (from light to dark red) indicates the average expression (log normalized counts) of a given marker within a given cell cluster.

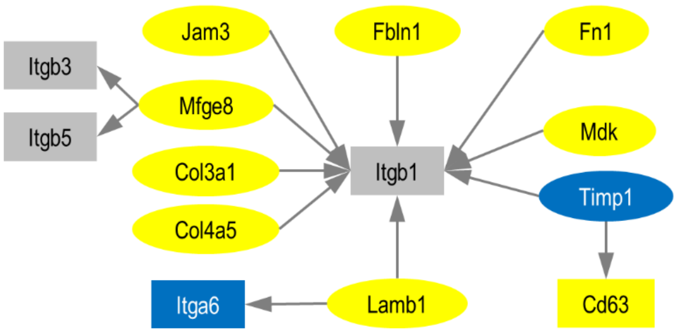


**Supplementary Figure 8. Selected ligand-receptor interactions between Sertoli cells (SC) and germ cells (GC).** Diagram of the ligand-receptor interactions between SC and GC, as revealed from the single cell transcriptomes. Each node of the network corresponds to a given gene encoding a ligand (oval shape) or a receptor (rectangle shape). Intercellular communication network among SC and GC were predicted using the CellTalkDB database, which contains literature-supported ligand-receptor pairs. Downregulated and upregulated genes are blue and yellow coloured, respectively. Grey boxes indicated receptors whose expression in not changed. The arrows point from the ligand to the receptors.
